# The neuroanatomical ultrastructure and function of a biological ring attractor

**DOI:** 10.1101/847152

**Authors:** Daniel B. Turner-Evans, Kristopher T. Jensen, Saba Ali, Tyler Paterson, Arlo Sheridan, Robert P. Ray, Tanya Wolff, Scott Lauritzen, Gerald M. Rubin, Davi Bock, Vivek Jayaraman

## Abstract

Neural representations of head direction have been discovered in many species. A large body of theoretical work has proposed that the dynamics associated with these representations is generated, maintained, and updated by recurrent network structures called ring attractors. We performed electron microscopy-based circuit reconstruction and RNA profiling of identified cell types in the heading direction system of *Drosophila melanogaster* to directly determine the underlying neural network. We identified network motifs that have been hypothesized to maintain the heading representation in darkness, update it when the animal turns, and tether it to visual cues. Functional studies provided additional support for the proposed roles of individual circuit elements. We also discovered recurrent connections between neuronal arbors with mixed pre- and post-synaptic specializations. Overall, our results confirm that the *Drosophila* heading direction network contains the core components of a ring attractor while also revealing unpredicted structural features that might enhance the network’s computational power.

## Introduction

Mammalian head-direction cells provide one of the clearest examples of an internal representation of an animal’s relationship to its surroundings (Taube et al., 1990a). The head-direction representation uses visual cues in the environment as a reference, but persists in darkness, where it is updated by self-motion cues (Taube et al., 1990b). This internal representation likely guides navigation behaviors. Indeed, perturbations to the system in rats induce errors in path-integration (Butler et al., 2017; Valerio and Taube, 2012). A large body of theoretical work has addressed conceptual questions about how such a representation of head direction might be generated and updated (Knierim and Zhang, 2012). These studies propose that the neuronal population dynamics associated with head-direction representations are maintained by network structures called ring attractors (Skaggs et al., 1995; Zhang, 1996). These recurrent networks, which are ideally suited to encoding a circular variable like head-direction, are often schematized as a ring of neurons whose connectivity depends on their directional tuning preferences (Figure 1A). In most model implementations, neurons with similar directional tuning excite each other and those with different tuning inhibit each other (Figure 1A), thereby enabling the generation of a stable pattern of localized activity in any part of the network (Amari, 1977; Ben-Yishai et al., 1995; Wu and Amari, 2005). Recurrent loops with a shift move the activity ‘bump’ around the ring as the animal turns (Figure 1B-D), and the compass-like representation is tethered to the animal’s surroundings through visual inputs, which are used as a reference to guide heading (Figure 1E,F). Experimental support for this general theoretical formulation has come from analyses of head-direction cell population activity under a variety of different conditions (Butler et al., 2017; Chaudhuri et al., 2019; Clark and Taube, 2012; Hargreaves et al., 2007; Muir et al., 2009; Peyrache et al., 2015; Taube et al., 1990b; Yoganarasimha et al., 2006). However, different network implementations of this general formulation make distinct assumptions about the connectivity of their constituent neurons (Ben-Yishai et al., 1995; Xie et al., 2002; Zhang, 1996) (for example, Figure 1A-F). Importantly, such assumptions, which determine exactly how the circuit functions, have been difficult to test in the large mammalian brain. Mammalian head-direction cells are distributed across many brain regions (Taube, 2007) and are as yet not well classified into types, making it challenging to identify and target them reliably.

**Figure 1:**
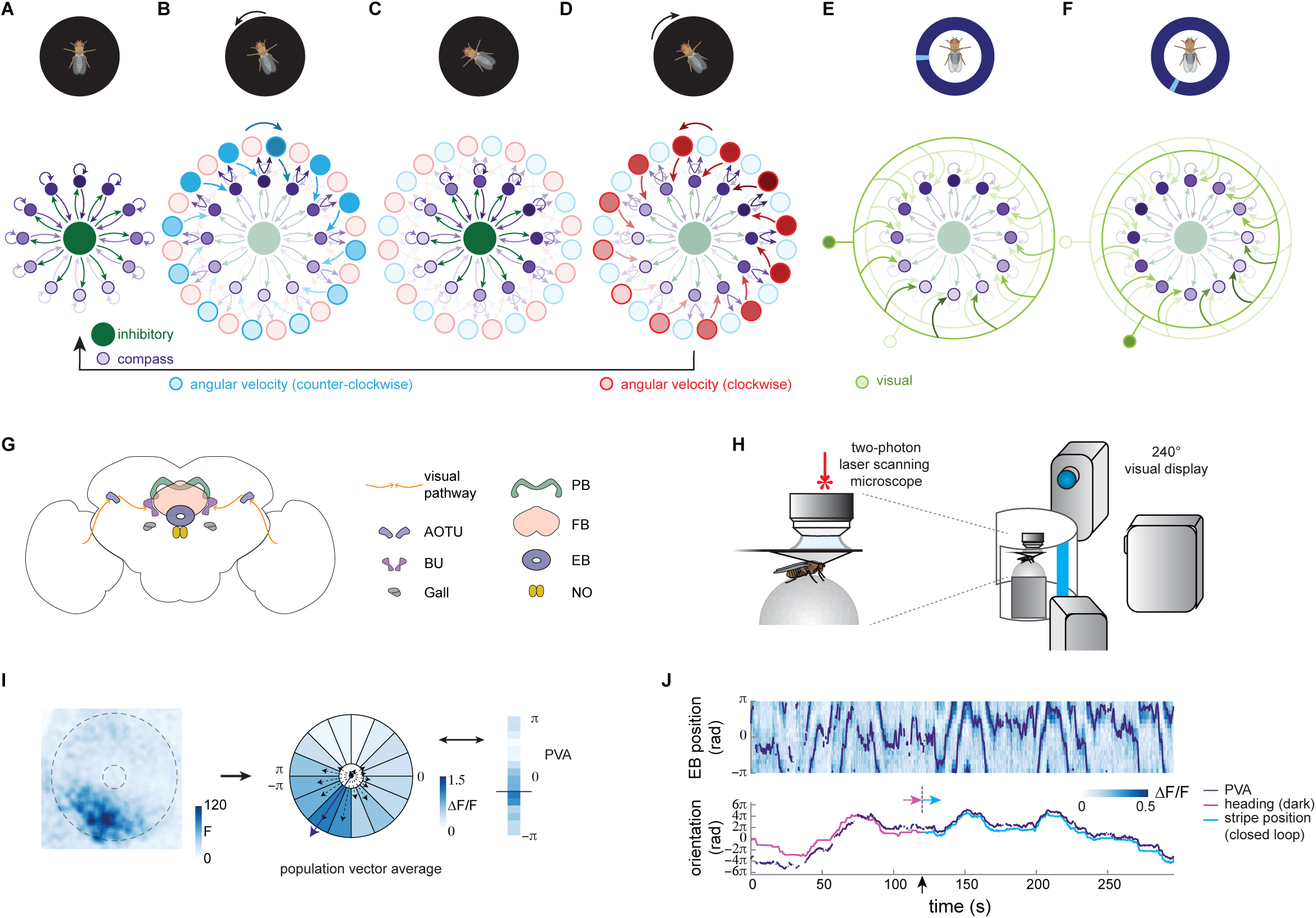
The fly neural compass as a ring attractor network. A-F: Function of the fly compass network schematized as a ring attractor in which the activity in each neuron is driven by excitatory angular velocity signals and inhibitory input from visual feature detectors. Darker shading indicates increased activity (the “bump”). The fly’s behavior and visual environment (shown in the cartoons on top) activate different subpopulations within the compass network. A. In the dark, local excitation (depicted for compactness as self-excitation) between compass neurons (purple) and uniform inhibition through inhibitory neurons (green, inside) leads to the formation of a ‘bump’ of activity that represents the fly’s heading direction (here “north”). B. Active compass neurons excite paired neurons that conjunctively tune to both heading and angular velocity. When the fly turns counterclockwise (CCW), the CCW angular velocity neurons (blue), in turn, excite compass neurons, shifting their activity clockwise (CW) (dark blue arrow, top). C. Shifted excitation from CCW angular velocity neurons moves the activity bump CW around the ring of compass neurons (dark purple). D. When the fly instead turns CW, the CW angular velocity neurons (red) shift the activity bump in the CCW direction (dark red arrow, top). E. Visual cues excite inhibitory visual feature detector neurons (green, outer ring) that suppress activity at all but a subset of compass positions. Here, a stripe at 9 o’clock activates a single visual neuron, which inhibits compass neuron activity everywhere except around 12 o’clock. F. When the visual cue (here a single stripe) moves, a new visual feature detector is excited. Here, a stripe at 7 o’clock excites a feature detector that permits compass neuron activity at 10 o’clock. G. A schematic of the fly central brain, focusing on a brain region called the central complex. The visual pathway into the central complex is shown with the orange arrows and progresses from the Anterior Optical Tubercle (AOTU) to the bulb (BU) and from the BU to the Ellipsoid Body (EB). Heading direction system neurons also arborize in the Gall, Protocerebral Bridge (PB), and Noduli (NO). The Fan-Shaped Body (FB) is the other central complex structure shown. H. Calcium activity of different neuron types in the compass network was measured in a head-fixed preparation. (left) Genetically expressed calcium indicators were imaged using two-photon laser scanning microscopy while a fly walked freely on an air-suspended Styrofoam ball. The ball rotation was monitored to track the fly’s orientation. (right) Visual stimuli (here, a 15° wide vertical stripe) were projected onto a 240° cylindrical screen with 3 projectors. Stimuli were either shown in open-loop (fly had no control of stimuli) or in one-dimensional closed-loop (the fly’s turns on the ball changed the angular position of the stimulus). I. (left) Fluorescence signal from GCaMP6f (here, in the E-PG neurons) showing the activity bump in the EB (outlined with the dotted lines). (right) The EB is divided into 16 equiangular sectors of interest. The change in fluorescence (ΔF/F) signal is then calculated in each of these regions. Δ/F is in turn used to calculate the Population Vector Average (PVA) to estimate the bump position. J. (top) An example of the E-PG EB activity over time. For the first 120 s, the fly is in darkness. For the final 150 s, a closed-loop stripe is shown. The PVA is overlaid in purple. (bottom) The unwrapped PVA (purple), the heading in the dark (magenta), and the stripe position (blue) are shown. The bump tracks the fly’s movements with accumulating error in darkness (before dotted line), and with minimal error in visual closed loop with a stripe stimulus (after dotted line).

An internal heading representation with similarities to the mammalian head-direction representation has also been discovered in the fly brain (Seelig and Jayaraman, 2015). Like the mammalian system, this heading representation tethers to visual surroundings and is maintained and updated in darkness. The fly’s experimental advantages —its small brain, identified neurons, genetic tools, and physiological tractability— make it an excellent system to assess the assumptions underlying the function of ring attractor networks (Figure 1A-F). Indeed, there is strong experimental and theoretical evidence from *Drosophila melanogaster* that the representation of heading is implemented by a ring attractor network (Green et al., 2017; Kim et al., 2017; Seelig and Jayaraman, 2015; Turner-Evans et al., 2017). A computational model of a ring attractor that assumes recurrent connectivity between heading neurons and neurons that encode both angular-velocity and heading (Figure 1B-D) has been shown to maintain an accurate representation of the fly’s heading when the circuit is driven by realistic velocity inputs (Turner-Evans et al., 2017). This model replicates the dynamics of the fly heading direction network in darkness. Other models have invoked plasticity between visual inputs and heading neurons to suggest how visual and angular velocity information might update the representation in a mutually consistent manner (Figure 1E,F) (Cope et al., 2017; Kim et al., 2019).

Importantly, however, these and other models of ring attractor networks in the fly have assumed the circuit’s connectivity based on relatively indirect evidence (Cope et al., 2017; Han et al., 2019; Kakaria and de Bivort, 2017; Kim et al., 2019; Kim et al., 2017; Su et al., 2017; Turner-Evans et al., 2017). For example, the location of pre- and post-synaptic arbors has been inferred from whether neural processes visible in light-microscopic images seem spine- or bouton-like in specific substructures (Hanesch et al., 1989; Lin et al., 2013; Wolff et al., 2015). The hypothesized connectivity of the circuit has then been derived from light-level overlap between the putatively pre- and post-synaptic arbors of neurons, in some cases further supported by <u>G</u>FP-reconstitution-<u>a</u>cross-<u>s</u>ynaptic-<u>p</u>artners (GRASP) (Xie et al., 2017) and trans-Tango (Omoto et al., 2018) experiments, although the reliability and accuracy of these methods to estimate pairwise connectivity is known to be limited (Lee et al., 2017; Talay et al., 2017). Similarly, measurements of functional connectivity by optogenetic stimulation of one neuron type and calcium imaging of another (Franconville et al., 2018) cannot conclusively establish the presence or absence of monosynaptic connectivity.

In the current study, we used reconstructions based on serial transmission electron microscopy (EM) (Zheng et al., 2018) to determine synaptic connectivity within the neural network underlying the fly’s heading representation. We supplemented this connectivity map with cell-type-specific RNA sequencing (RNA-Seq) and fluorescence in situ hybridization (FISH), which allowed us to characterize the expression profiles of the key cellular components of the ring attractor network. We then used this integrated information to quantitatively assess the role of each of the constituent cell types in the ring attractor’s dynamics. We found that the structure of the heading direction network contains motifs similar to those proposed in theoretical models. These motifs were hypothesized to maintain the heading representation activity and update it both in the dark and when visual features are present. We tested these ideas using targeted two-photon calcium imaging and thermogenetics of the constituent neuron types in behaving, head-fixed *Drosophila*. We also found that many neurons have mixed pre- and post-synaptic specializations within their innervations to single brain structures, creating “hyper-local” recurrent loops that may allow local computations to supplement the role of recurrence at the network level. Moreover, while many implementations of ring attractor networks rely on distinct units that provide local excitation and long-range inhibition to shape activity into one stable bump, consistent with our results, we found apparent redundancy in these structural elements. Both local excitation and long-range inhibition appear to be carried out by multiple classes of neurons. Taken together, our results provide new structural and functional insights into how a small biological ring attractor network allows an animal to maintain an accurate internal sense of direction.

## Results

The fly’s heading is topographically represented in the toroid-shaped ellipsoid body (EB), a structure within a brain region called the central complex (Figure 1G). The representation manifests as a localized ‘bump’ of population activity in the so-called E-PG neurons (see Methods for notes on nomenclature) (Seelig and Jayaraman, 2015). Neurons in the E-PG population together tile the EB, with each neuron’s arbors occupying a single ∼22.5° wedge of the EB (Wolff et al., 2015). E-PG neurons in nearby wedges are tuned to similar heading directions and those in angularly distant wedges to different heading directions, resulting in a complete topographical representation across the circumference of the EB. To observe E-PG activity, we placed tethered, walking flies on a ball inside a visual virtual reality (VR) setup (Figure 1H). The setup permitted the fly to control its angular orientation relative to visual cues presented (see Methods). We exposed the fly’s brain and performed two-photon laser scanning microscopy using the genetically encoded calcium indicator, GCaMP6f (Chen et al., 2013) (see Methods), which we expressed selectively in E-PG neurons (Figure 1I). As in previous studies, we characterized E-PG population dynamics by computing and tracking the population vector average of E-PG activity (Figure 1I) in flies walking in darkness and in closed-loop VR in simple, single-stripe visual environments (Fisher et al., 2019; Green et al., 2017; Green et al., 2019; Seelig and Jayaraman, 2015; Turner-Evans et al., 2017). The E-PG bump tracked the fly’s heading in darkness and, more reliably, in our closed loop visual VR conditions (Figure 1J, Figure S1A,B). We also quantified the stability of the heading representation over time, confirming that the bump reliably tracked the heading over a 4 s sliding window across the trial (Figure S1C-E, see Methods). Having confirmed that the heading representation accurately integrates the fly’s heading, we sought to characterize the cell types and network motifs that together generate the E-PG bump and its ring-attractor-like dynamics.

### Cellular- and synaptic-resolution network motifs underlying the ring attractor network

We first focused on uncovering —at cellular and synaptic resolution— the motifs that together constitute the core of the ring attractor network (Figure 1A-F). For this, we relied on a recently released synaptic-resolution whole-brain EM dataset (Zheng et al., 2018) (Figure 2A). Note that the resolution of our data did not permit the identification of electrical synapses. We reconstructed several neurons of each of the cell types that arborize within two of the brain regions where the heading representation has been studied previously in the fly: the EB and protocerebral bridge (PB) (Figure S2) (Green et al., 2017; Seelig and Jayaraman, 2015; Turner-Evans et al., 2017). We reconstructed selected neurons to morphological completion in both of these brain areas and annotated all synaptic contacts between the reconstructed neurons (Figure 2B, see Methods).

**Figure 2:**
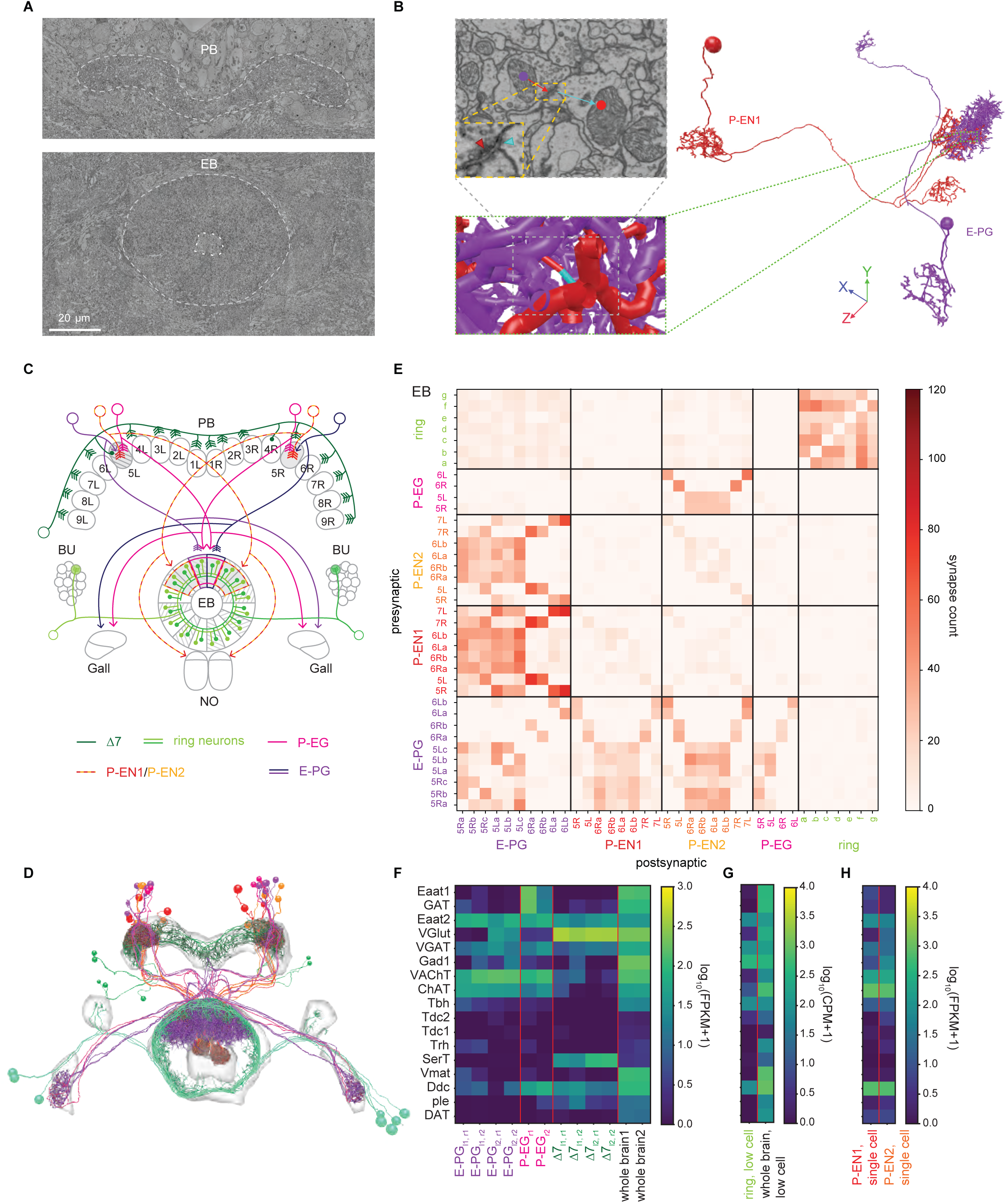
Electron-microscopic reconstruction and. A. Cross-sections of FAFB electron microscopic volume showing the PB (top) and the EB (bottom). B. An example synapse between an E-PG neuron (purple) and a P-EN1 neuron (red) in the ellipsoid body. (top left) Electron microscopy view. The inset shows the t-bar (red carat) and post-synaptic density (blue carat). (bottom left) Projected view of the 3D skeleton showing the synapse location between skeletons. (right) Complete skeletons of the two neurons showing the synapse location. C. Schematic showing examples of neurons that are thought to play a role in the heading direction ring attractor. Neurons are color coded according to their putative roles in the attractor as shown in Figure 1A-F. Individual neurons are labeled according to where they arborize in the PB, following the numbering scheme here. Columnar neurons (E-PGs, P-ENs, and P-EGs) link subregions of the EB and PB while tangential neurons (Δ7s and ring neurons) arborize throughout either the PB or EB. D. Skeletons of the subset of neurons that were manually traced in the FAFB dataset. Synapses were manually annotated but are not shown. The light grey regions show the outlines of the central brain neuropil. The neurons are colored as in Figure 2C. E. Connectivity matrix for the traced neurons in the ellipsoid body. F-H. RNA sequencing was performed on the cell classes identified in Figure 2C. Genes related to transmitters are shown here. Three types of sequencing pipelines were used: bulk (F), low cell (G), and single cell (H) (See Methods). F. Bulk cell sequencing of E-PG, P-EG, and Δ7 neurons along with whole brain samples. l1 and l2 refer to distinct genetic lines while r1 and r2 refer to distinct technical replicates. G. Low cell sequencing was performed on the ring neurons and on whole brain samples. The mean signal across biological replicates (7 for the ring neuron line and 8 for the whole brain material) is shown here. H. Single cell sequencing was performed on the P-EN1 and P-EN2 genetic lines, which contained additional cell types. The mean signal across the samples is shown here.

We focused on all the known ‘columnar’ neuron types (E-PG, P-EN1, P-EN2, P-EG) that link specific EB wedges with corresponding columns in the PB, and on a subset of known ‘tangential neurons’ (ring and Δ7 neurons) within the specific sub-regions of the EB and PB that were innervated by the columnar neurons (Figure 2C-D; also see note on nomenclature in Methods) (Hanesch et al., 1989; Lin et al., 2013; Wolff et al., 2015; Wolff and Rubin, 2018). Based on the stereotyped morphologies and tiled projection patterns of each of these cell types within the EB and PB, we expect that the connectivity patterns we uncovered within these selected sub-regions will generalize to the other wedges and columns in the EB and PB. We marked every pre- and post-synaptic site in each neuron that we traced and constructed connectivity matrices for each region (Figure 2E). While this approach allowed us to identify how often these chosen neurons synapsed onto one another, it still left many partners unidentified. To check the completeness of our synapse partner identification, we therefore looked at the percent of inputs and outputs that were labeled (See Methods for synapse distribution protocol, Figure S3). We could identify approximately 60% of the total inputs in the PB and EB, approximately 40% of the inputs in the NO, and approximately 40% of all outputs across regions. The unidentified connections are to neurons that we did not reconstruct.

Next, to determine the signs of the connections between identified partners —that is, whether synaptic connections are excitatory or inhibitory— we performed RNA-Seq on each cell type that we could target specifically with GAL4 driver lines (Figure 2F-H; see Methods) (Aso et al., 2019; Davis et al., 2019). For some cell types, we also used FISH in parallel to confirm the likely neurotransmitter (Long et al., 2017; Meissner et al., 2019). Most cell types appear to express only one neurotransmitter and likely express receptors for most, if not all, neurotransmitters (Davis et al., 2019). Both receptor and neurotransmitter identity are needed to infer the likely sign of a synaptic connection between two neurons. E-PGs, P-EGs, P-EN1s, and P-EN2s all express VAChT and ChAT identifying them as cholinergic (Figure 2F,H). The subclass of ring neurons that we targeted, likely visual neurons, expressed Gad1, the enzyme that produces GABA (adjusted p=6.25E-5 with respect to the whole brain data), identifying them as GABAergic (Figure 2G, see Supplemental Information for neuropeptide signaling) (Enell et al., 2007; Homberg et al., 1999; Kahsai et al., 2012; Martin-Pena et al., 2014; Zhang et al., 2013). The Δ7s expressed VGlut, marking them as glutamatergic (Figure 2F, adjusted p=3.08E-2) (Daniels et al., 2008). Receptor identity will be discussed in the following sections. Next, having measured the synaptic connectivity matrix for the key components of the EB-PB heading direction network, and having gained some insight into their synaptic and cellular properties from RNA-Seq and FISH, we asked how these different cell types and motifs might underlie specific properties of the ring attractor network.

### E-PG neuron output is essential to the maintenance of the heading representation

The fly ring attractor network is thought to enable the formation of a single, stable bump of activity through local excitation and near-uniform inhibition (Figure 3A) (Kim et al., 2017). Local excitation reinforces activity at the bump location while long-range inhibition suppresses activity in neurons with different heading tuning. In prior work (Turner-Evans et al., 2017), we had suggested that the E-PG neurons are the center of the ring attractor network, and that their role in driving excitatory recurrent loops was essential to the generation of a stable heading representation. We further suggested that the Δ7 neurons might provide the necessary long-range inhibition. Here, we test this assumption at the level of connectivity and functional activity in the E-PG and Δ7 neurons (Figure 3B,C).

**Figure 3:**
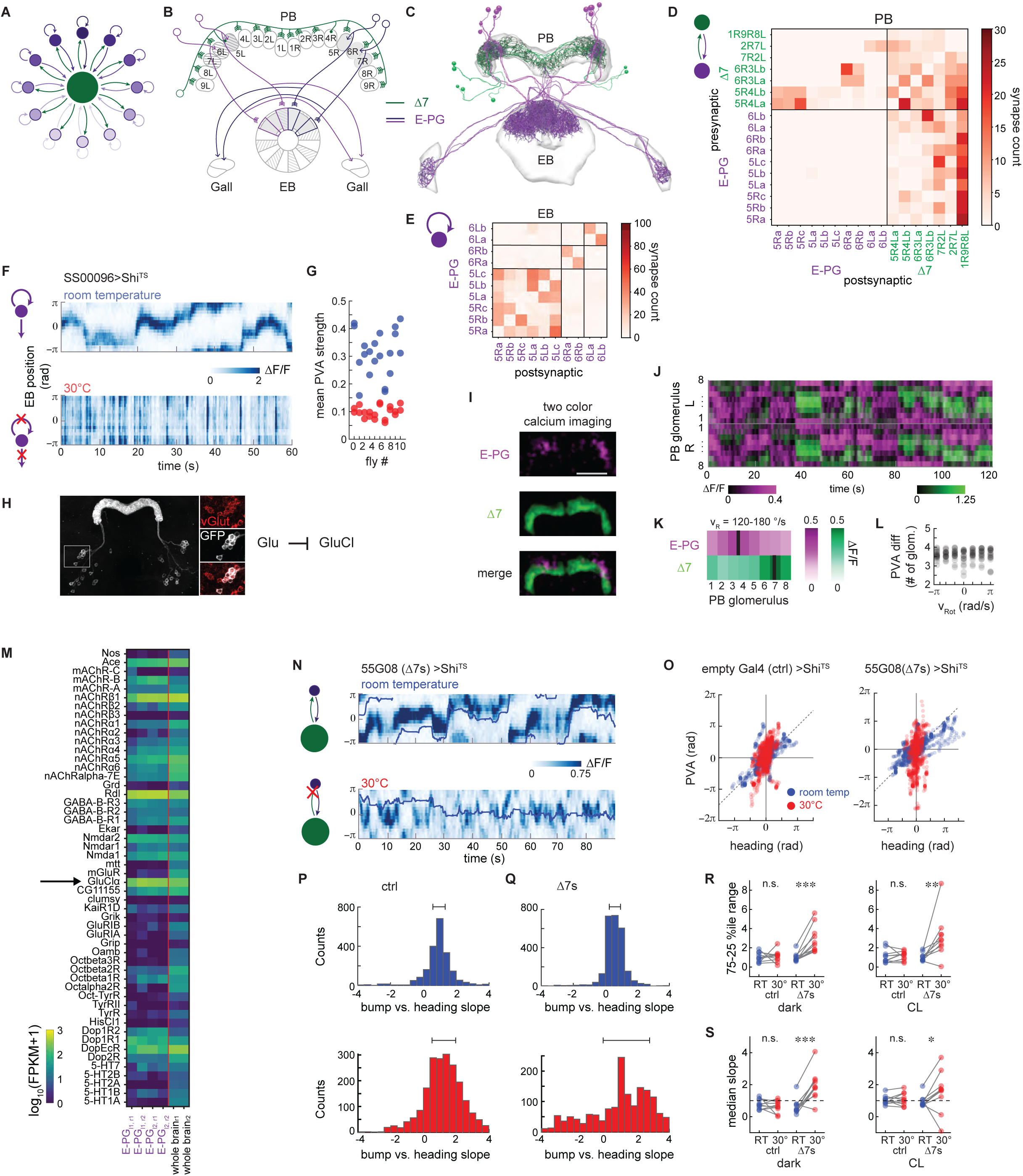
Local excitation from E-PGs and long-range inhibition from Δ7s are required to stabilize the E-PG activity bump. A. Local excitation and long-range inhibition lead to the formation of one bump of activity (dark purple). B. Schematic depicting a subset of the E-PG and Δ7 neurons corresponding to those traced (Figure 3C). Grey shading shows areas of PB and EB in which the neurons were traced. Note that Δ7s were also partially traced in other PB glomeruli to enable identification. C. Putative excitatory (purple, E-PGs) and inhibitory (green, Δ7s) neurons were traced and their synapses onto one another annotated. D. Connectivity matrix within the protocerebral bridge (PB) for the E-PGs Δ7s. The Δ7s synapse onto the subset of E-PGs within one glomerulus in the PB (5R→, 6R 6R, 5R → upper left quadrant) and receive broad input from the E-PGs away from that region (last column, lower right quadrant). The Δ7s synapse onto one another (upper right quadrant). E. Connectivity matrix within the ellipsoid body (EB). The E-PGs within a wedge are densely interconnected. Some E-PGs also make a few synapses onto their counterparts from neighboring wedges. F. Calcium activity in the ellipsoid body of the E-PGs as recorded with GCaMP6f for a fly expressing shi^TS^ in the E-PGs (SS00096). The fly is walking in darkness. At room temperature (top), a bump of activity is clearly visible. At the restrictive temperature (bottom), when synaptic vesicle reuptake is blocked within the E-PGs, the bump is no longer visible. Instead, E-PGs across the entire ellipsoid body flash off and on. G. The PVA strength as a function of temperature across 10 flies. Restrictive temperature trials are shown in red, while permissive temperature trials are shown in blue. The PVA strength is the mean resultant vector length across the 16 ROIs that span the ellipsoid body (see Figure 1I). A PVA strength of 1 indicates that all of the activity is in one ROI, while a PVA strength of 0 indicates that the activity is spread equally across all ROIs. The average PVA strength for a given trial is shown. H. Fluorescent *in situ* hybridization (FISH) of vGlut showed consistent overlap between GFP labeled Δ7s (white) and vGlut positive cells (red) (3/3 brains for the SS30295 genetic line and 2/2 brains for the SS02238 genetic line). (right) Glutamate release can lead to hyperpolarization of a downstream partner through GluCl channels. I. Two-color calcium imaging was performed on the E-PGs (magenta) and the Δ7s (green) to record the simultaneous activity of the two populations. Here, GCaMP6f was expressed in the Δ7s jRGECO1a was expressed in the E-PGs. J. The PB activity of both populations over time for the fly shown in I. The right and left PB were each divided into eight regions of interest (ROI), each corresponding to one glomerulus. The change in fluorescence intensity, ΔF/F, was then calculated for each ROI over time. K. At each point in time, the red and green activity profiles were circularly shifted by the same number of ROIs in order to consistently place the peak of red activity at ROI #4. The mean registered activity in the right half of the PB across all time points when the fly in H was rotating at 120-180 °/s is shown. L. The offset between peak E-PG and Δ7 activity as a function of rotational velocity, v_RoT_. Data from 10 flies is shown, with both the right and left PB included. In 5 flies, Rot GCaMP6f was expressed in the Δ7s and jRGECO1a was expressed in the E-PGs, while 5 flies had the reverse indicator expression pairing. M. Expression of genes linked to receptors as compiled from RNA sequencing of the E-PGs and whole brain tissue (see Methods). mRNA expression of the GluClα subunit is highlighted. N. E-PG calcium activity in the ellipsoid body recorded with GCaMP6f in flies expressing shi^TS^ in the Δ7s (line 55G08) at permissive (top) and restrictive temperatures (bottom). The fly was presented with a stripe in closed loop over the course of the trial, and its position over time is indicated by the blue line. O. For 4 s sliding windows (see Methods), the fly’s position (heading) is plotted vs. the bump position (PVA). Blue dots indicate trials at the permissive temperature and red dots indicate trials at the restrictive temperature. (left) Heading vs. PVA for an example control fly. (right) Heading vs. PVA for an example fly expressing shi^TS^ in the 7s. Δ P. The slope of the points in 4 s sliding windows. The two plots shown here are for flies expressing GCaMP6f in the E-PGs and shi^TS^ in empty Gal4 controls. The top plot shows a histogram of the slopes across two room temperature trials. The bottom plot shows a histogram of the slopes across three high temperature trials. The 25^th^ and 75th percentiles are indicated above. Q. As in Figure 3P, but now for flies expressing shi^TS^ in the 7s. Δ R. The range between the 25th and 75th percentiles, as obtained from the histogram of slopes, for the flies expressing shi^TS^ in empty Gal4 (ctrl) and the Δ7s. (left) The percentile range when the flies are in the dark. (right) The percentile range when the flies are tracking a closed loop stripe. Trials at the permissive temperature are in blue; trials at the restrictive temperature are in red. From left to right, p = 0.81, 5E-4, 0.80, and 9.7E-3 (paired t-test). S. The median slope for the control flies in the dark (left) and with a stripe (right) for the room temperature trials (blue) and high temperature trials (red). From left to right, p = 0.43, 3.5E-4, 0.61, and 0.13 (paired t-test).

Our data revealed that E-PG neurons are cholinergic (likely excitatory, Figure 2F), and are the sole columnar neuron type that both receives synaptic input in the EB and provides a significant number of synaptic outputs to neurons in the PB (Figure 3D), consistent with their proposed role as the primary, excitatory neural population in the fly ring attractor network. In addition, examining the connectivity between E-PGs within the EB (Figure 3E), we found that the neurons within a given wedge are not purely post-synaptic (Lin et al., 2013), but rather make synapses onto one another, consistent with the locally excitatory connectivity predicted by many ring attractor models (Kim et al., 2017). However, a key question about the E-PG neurons is whether their compass-like activity is entirely derived from their inputs or whether they are themselves required to generate these dynamics. If the heading representation is inherited from other cell types that provide input to the E-PG population, blocking E-PG outputs should leave their heading tuning unimpaired. By contrast, if E-PGs themselves are an important hub in the heading direction network, as has been strongly suggested previously (Kim et al., 2017; Turner-Evans et al., 2017), blocking the E-PG outputs should destroy their heading tuning. To answer this question, we expressed both GCaMP6f and the temperature-sensitive mutation of the *Drosophila* dynamin orthologue, shibire^ts1^ (shi^ts^)(Kitamoto, 2002), which blocks vesicle endocytosis and thus synaptic transmission at elevated temperatures, in the E-PGs and measured their Ca^2+^ activity at both permissive and restrictive temperatures. At permissive temperatures, one bump of activity was clearly visible and tracked the animal’s movements (Figure 3F, top). In contrast, at higher temperatures, the activity in the entire EB instead rapidly increased when the animal turned and decreased shortly thereafter (Figure 3F bottom, Figure S4A-D; note that the E-PG bump tracks angular orientation in flies expressing empty-Gal4 and shi^ts^ (Figure 5G, I, J, Figure S4I)). The magnitude of the population vector average (PVA, a measure of the degree of localization of E-PG activity) dropped drastically across flies at the restrictive temperature (Figure 3G), suggesting that the E-PGs are in fact required to generate compass-like activity dynamics.

### The Δ7 neurons stabilize the heading representation by inhibiting E-PG neurons

Long-range inhibition has been a necessary stabilizing feature of all ring attractor networks, including those proposed for the fly heading direction network (Kakaria and de Bivort, 2017; Kim et al., 2017; Turner-Evans et al., 2017). A prominent candidate for this inhibition has been the Δ7 class of neurons (Heinze and Homberg, 2007; Wolff et al., 2015), which have been shown to functionally inhibit the E-PG population in the PB (Franconville et al., 2018). We first asked if this functional inhibition arises from direct synapses between individual Δ7 neurons onto E-PG neurons. Our EM reconstructions confirmed that E-PGs indeed receive synapses from Δ7 neurons (Figure 3D). Our FISH (Figure 3H) and RNA-Seq (Figure 2F) analysis both confirmed previous results suggesting that the Δ7s were glutamatergic (Daniels et al., 2008). In the fly, glutamate can bind to many different types of receptors on downstream partners. NMDA and some mGluR receptors are believed to lead to excitatory responses, while GluClα receptors lead to inhibitory responses (Liu and Wilson, 2013; Xia et al., 2005). The RNA-Seq data revealed GluClα mRNA expression in the E-PG neurons (Figure 3M, adjusted p=0.386), raising the possibity that GluC1Δ channels underlie their inhibition by the Δ7 population (Franconville et al., 2018). Consistent with the columnar specificity of their pre- and post-synaptic specializations (Heinze and Homberg, 2007; Wolff et al., 2015), we found that Δ7 neurons synapse selectively onto only the subset of E-PGs within one column in the PB (Figure 3D, upper left quadrant of the matrix) and receive broad input from the E-PGs away from that region (Figure 3D, lower right quadrant). Overall, the connectivity and RNA-Seq data are consistent with Δ7s putatively inhibiting those E-PGs that correspond to angular orientations distant from the current orientation.

We also discovered that the Δ7s synapse strongly onto one another (Figure 3D, upper right quadrant), indicating that these neurons may play a much more nuanced computational role than their light-level anatomy suggests. Unfortunately, our RNA-Seq results for the receptors expressed by Δ7 neurons were inconclusive, with relatively weak signatures of both GluClα and expressed by mGluR.

In many ring attractor models, a single activity bump is ensured by assuming that active head-direction neurons (in our case, E-PG neurons) inhibit others with distant angular orientation tuning preferences. If the Δ7 population performs such a role, we would expect these neurons to be most active in columns in which the E-PGs are inactive and vice versa. We performed simultaneous two-color imaging of the Ca^2+^ activity of both populations of neurons, labeling one with the green indicator GCaMP6f and one with a red indicator, jRGECO1a (Figure 3I). As predicted, we found that a bump of Δ7 activity peaked in exactly the opposite angular orientation as the E-PG bump, so that the spatial locations of their respective activity bumps are offset 180° with respect to one another (Figures 3I3I-L, Figure S4H). Taken together, the offset between the Δ7 and E-PG bumps, along with the morphology, expression profile, and connectivity of the Δ7 neurons, suggests that the Δ7 neurons provide long-range inhibition that helps maintain a single stable E-PG bump. We used shi^ts^ to perform perturbation experiments to ask if the Δ7s are, in fact, the only source of inhibition in the heading perturbation experiments to ask if the direction network. Specifically, we expressed GCaMP6f in the E-PGs and shi^ts^ in the Δ7s, and measured E-PG Ca^2+^ activity at both permissive and restrictive temperatures. If the Δ7s are the main source of inhibition in the ring attractor network, we expect that blocking their outputs would cause the overall activity of the E-PGs to increase and the bump shape to be altered, potentially even leading to multiple bumps. However, this is not what we observed. At the restrictive temperature, the bump was visible as before, but now moved erratically in response to the fly’s movements (Figure 3N bottom). Further, although the bump amplitude was lower in these flies as compared to control flies, the bump width remained the same (Figure S4J,K). For control flies, at both low and high temperatures, the bump accurately tracked the stripe (Figure 3O left, P, Figure S4I; see Methods). The flies expressing shi^TS^ in the Δ7s have similar bump dynamics at permissive temperatures (Figure 3N top, blue dots in Figure 3O, right), but are less consistent at the restrictive temperatures (Figure 3Q-S; see Methods). This quantification confirmed that the bump no longer reliably tracks the fly’s movements when Δ7s are inhibited.

These observations suggest that other, as yet unidentified, sources of inhibition must act to shape the E-PG activity into one bump while the Δ7 neurons contribute to stabilizing the bump’s movements. P-EN1 neurons recurrently connected to E-PGs update the compass during turns Ring attractor theories typically invoke a distinct class of neurons that are tuned to both angular velocity and heading to update the heading representation as the animal turns in darkness (Figure 1B-D) (Skaggs et al., 1995; Xie et al., 2002; Zhang, 1996). Previous studies provided strong evidence that P-EN1 neurons perform this role in the fly (Green et al., 2017; Turner-Evans et al., 2017). P-EN1 neurons are believed to shift the heading representation around the ring through clockwise and counter-clockwise anatomical offsets (Figure 4A). Light-level characterization and functional connectivity experiments have suggested that P-EN1 neurons receive input from the E-PGs in the PB (Lin et al., 2013; Turner-Evans et al., 2017; Wolff et al., 2015). These neurons then send their outputs to E-PGs in the EB in wedges that are adjacent to those innervated by the E-PGs that provided the same PEN1s with input (Figure 4B) (Green et al., 2017; Turner-Evans et al., 2017; Wolff et al., 2015). We examined whether these connections are direct and synaptic at the level of EM (Figure 4C). In the PB, the E-PGs indeed directly contact the P-EN1s as predicted (Figure 4D). They also likely inhibit P-EN1s in distant PB glomeruli (and therefore distant angular orientation tuning) through the Δ7 population (Figure S4M). In turn, the P-EN1s connect most strongly to anatomically shifted E-PGs in the EB (Figure 4E, top left quadrant of the matrix). Interestingly, while there are varying numbers of P-EN1s that link one glomerulus in the PB to one sector in the EB (1 for some columns, 2 for others), the total number of synapses between the P-EN1s and E-PGs in a given sector remains roughly constant (Figure 4F).

**Figure 4:**
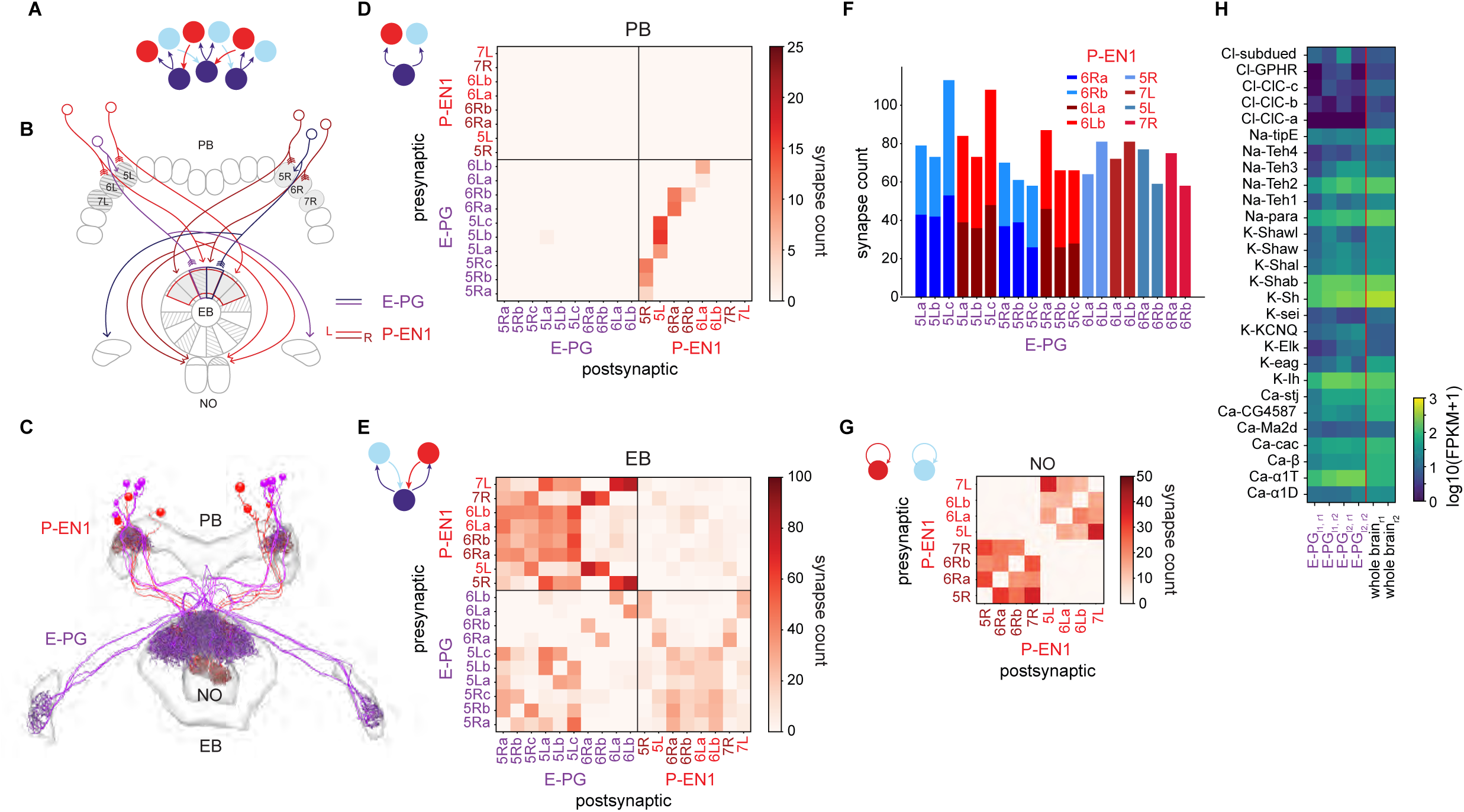
Recurrent loop between E-PGs and P-EN1s that update bump position when the animal turns is angularly shifted in the EB. A. Neurons that conjointly encode the fly’s angular velocity and heading (red, light blue) move the bump (purple) around the ring (see Figure 1B). B. Schematic displaying the anatomically-shifted recurrent loop between sample E-PGs and P-EN1s. EB wedges and PB glomeruli selected for E-PG and P-EN1 reconstruction (Figure 4C) are in grey. C. E-PG and P-EN1 neurons were manually traced and their synapses onto each other annotated. D. Connectivity matrix between the E-PGs and the P-EN1s in the PB. The E-PGs synapse onto the P-EN1s that arborize in the same glomerulus (lower right quadrant) E. Connectivity matrix between the E-PGs and the P-EN1s in the EB. The shifted P-EN1s synapse onto the E-PGs (top left quadrant, e.g. P-EN1 6R→E-PG 5L). Note that the E-→PGs also synapse onto the P-EN1s in the EB (bottom right quadrant). F. The total number of synapses from P-EN1s onto individual E-PGs. Although some individual E-PGs receive inputs from single P-EN1s and others from two P-EN1s, the total number of synapses is approximately maintained (p = 0.295, two sample t-test, n = 12 samples for two P-EN1s, n = 8 samples for one P-EN1s). G. Connectivity matrix between P-EN1s in the NO. Note that P-EN1s from the same side of the PB are heavily interconnected in the NO. H. mRNA expression of voltage-gated channels in the E-PGs.

We assessed the functional significance of these localized and precisely structured recurrent loops involving the E-PG and P-EN1 neurons by examining bump dynamics when E-PG outputs were blocked using shi^ts^. As mentioned in a previous section, this manipulation resulted in the entire E-PG population becoming active whenever the animal turned (Figure 3F, Figure S4C,D), likely as a result of uniform P-EN1 inputs that were no longer sharpened in their heading tuning by excitation from the E-PGs or by structured inhibition from the Δ7 population (Figure S4M). In some flies, this excitation was uniform across the EB, but in other flies one region was more excited than others, or different regions were more strongly excited for different turn directions, suggesting that P-EN1 or other inputs may be heterogeneous in these flies (Figure S4G, flies 2-4).

EM reconstruction and synapse annotation also revealed unexpected connections from the E-PGs back to the P-ENs within the EB (Figure 4E, lower right quadrant). Thus, within the EB, these two classes of neurons form “hyper-local” feedback loops, with pre- and post-synaptic sites located within the same processes in the EB. This may contribute to the persistence of the heading representation in the absence of sensory input (other potential contributing mechanisms are suggested below and in Discussion). Persistent activity in the E-PGs could also be sustained by voltage-gated channels and prevented from running away by inhibitory autoreceptors. Indeed, RNA-Seq of E-PGs reveals expression of several voltage-gated channel-linked genes (Figure 4H) and mAChR-B mRNA (Figure 3M, adjusted p=7.42E-3). The mammalian homologue of mAChR-B, M4, can act as an inhibitory autoreceptor for acetylcholine (Zhang et al., 2002).

The P-EN1s also arborize within a third brain structure, the noduli (NO). Within this structure, the P-EN1s synapse onto one another (Figure 4G), but the function of this local recurrence is not yet clear.

In summary, consistent with previous functional observations (Green et al., 2017; Turner-Evans et al., 2017), the P-EN1 population’s connectivity is consistent with their proposed role in updating the bump position. Further, unexpected local connections between P-EN1s and E-PGs in the EB point to its potential involvement in regulating sustained E-PG activity when the fly is standing still.

### Ring neurons tether the compass to visual scenes by selectively inhibiting E-PG neurons

Ring attractor models of head direction networks provide a mechanism for angular integration in the absence of visual cues, but such models (and the real networks that they model) accumulate error over time in darkness. Indeed, the fly compass is more accurate at representing the fly’s heading direction in visual closed-loop conditions than in darkness (Figure 1J, Figure S1A,B) (Fisher et al., 2019; Seelig and Jayaraman, 2015). Most head direction network models thus incorporate visual inputs (Figure 5A), and many hypothesize that the synapses between these inputs and compass neurons should be plastic, allowing visual cues to flexibly map onto and thereby tether the heading representation (Figure 1E-F) (Cope et al., 2017; Kim et al., 2019; Knierim et al., 1998; Ocko et al., 2018; Page and Jeffery, 2018).

**Figure 5:**
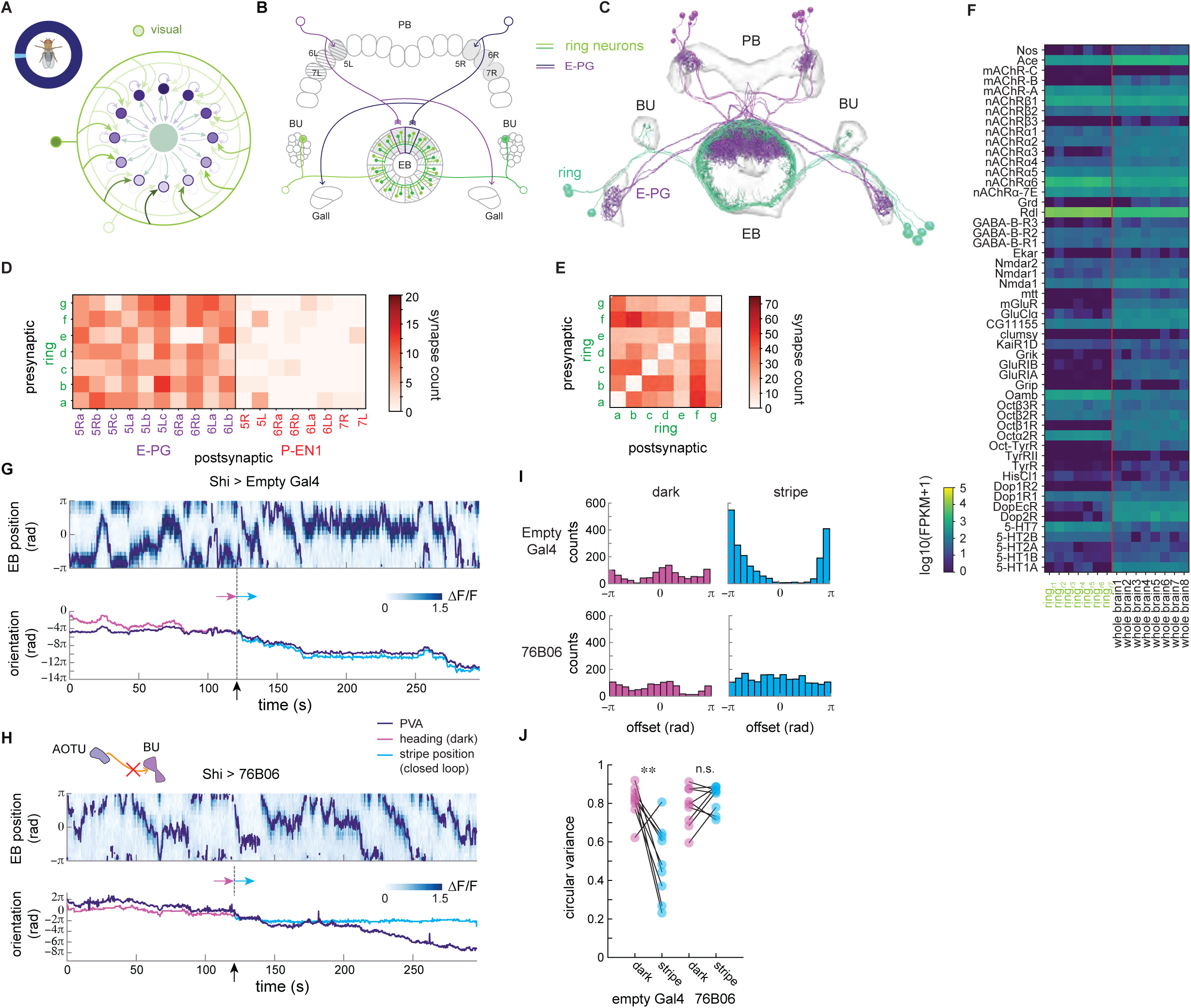
Mutually inhibitory R4d ring neurons synapse onto the E-PGs, tethering the bump to visual cues. A. Visual inputs (light green) map the visual scene onto the compass bump (dark purple) (see Figure 1E,F for details). B. Schematic displaying ring neurons making synaptic contacts onto E-PGs in the EB. EB wedges and PB glomeruli selected for E-PG reconstruction (Figure 5C) are in grey. C. R4d neurons (light green) were traced and their synapses onto each other and onto E-PG neurons (purple) were annotated. D. Connectivity matrix between the E-PGs, P-EN1s, and ring neurons in the EB. E. Connectivity matrix between ring neurons in the EB. F. mRNA expression of receptor-linked genes for the ring neurons and whole brain tissue. Subscripts r1-r7 refer to biological replicates, each of which uses the same genetic line to target the R4d ring neurons. G. An example of E-PG EB activity over time for control flies at the restrictive temperature (top). For the first 120 s, the fly is in darkness (heading shown in pink). For the final 150 s, the fly sees a stripe in closed loop (stripe position shown in light blue). The PVA is overlaid in purple. (bottom) Unwrapped PVA, heading in the dark, and stripe position over time. H. An example of E-PG EB activity for flies expressing shi^TS^ in the TuBu neurons at the restrictive temperature. Compare E-PG bump locking to visual cue movement when the fly is walking in closed loop with a stripe against that seen at permissive temperature (Figure 5G). I. The offset between PVA and heading is calculated at each time point for the trials shown in Figure 5G and Figure 5H, binned, and plotted here. J. The circular variance for the offset distribution across flies in the dark and when a closed-loop stripe is present (p = 4.9E-3, 0.23 (paired t-test)).

In *Drosophila*, this tethering is thought to come from a large population of inhibitory ring neurons (Figure 5B) (Hanesch et al., 1989; Homberg et al., 2018; Lin et al., 2013; Omoto et al., 2018; Xie et al., 2017; Young and Armstrong, 2010), an assumption with recent experimental support (Fisher et al., 2019). A potential experimental complication is that there are many types of ring neurons, only some of which are visually tuned (Homberg et al., 2011; Omoto et al., 2017; Omoto et al., 2018; Seelig and Jayaraman, 2013; Shiozaki and Kazama, 2017; Sun et al., 2017). To identify appropriate visual ring neurons to reconstruct in the EM dataset, we performed stochastic labeling of broad ring neuron lines using the FLP-out technique (Figure S5A) (Golic and Lindquist, 1989) (see Methods). This technique allows for individual neurons to be tagged, and here we used it to express GCaMP6f in single neurons in order to monitor their Ca^2+^ activity. This procedure labeled a wide array of morphologically distinct ring neurons, and we measured the activity of each type in response to open-loop rotating stripes (Figure S5A). Consistent with expectations from previous studies (Fisher et al., 2019; Omoto et al., 2017; Seelig and Jayaraman, 2013; Shiozaki and Kazama, 2017; Sun et al., 2017), R4d ring neurons consistently showed strong, ipsilateral responses to rotating stripes presented in open loop in contrast to several other types of ring neurons (Figure S5A). We thus focused our EM reconstruction efforts on the R4d class (Figure 5B,C), while noting that the R2 ring neurons are also known to be visually responsive (Fisher et al., 2019; Omoto et al., 2017; Seelig and Jayaraman, 2013).

We found that R4d neurons selectively target the E-PGs over the P-EN1s (Figure 5D), consistent with most models of visually tethered ring attractor networks. Insect compass models that incorporate visual input (Cope et al., 2017; Kim et al., 2019) suggest that ring neurons that respond to visual cues at specific orientations should more strongly inhibit E-PGs away from the desired bump location, creating a less-inhibited trough for the bump of activity to settle into for those orientations (Fisher et al., 2019). Such a trough is not visible in the synaptic counts of the ring neurons we reconstructed. Instead, the number of synapses between the ring neurons and the E-PGs is relatively constant across the sectors of the EB that we reconstructed. The trough may indeed be represented in synapse numbers, but it may not be visible across the limited section of the EB that we traced, or it may instead be represented in the strengths of the synapses but not in their numbers.

A surprise in the EM reconstructions was how heavily R4d ring neurons synapse onto each other (Figure 5E). RNA-Seq of these neurons revealed relatively high expression levels of GABA related receptor genes as compared to the whole brain data (Figure 5F, adjusted p=1.86E-19 for Rdl), suggesting that ring neurons are inhibited by GABA and therefore by each other. This hypothesis is consistent with recent observations that combined optogenetic stimulation of individual R2 ring neurons with whole-cell patch clamp electrophysiological recordings of other R2 neurons (Isaacman-Beck et al., 2019). When we compared the calcium activity of the R4d population in darkness to the activity seen in the presence of a single bright stripe moving in open loop, we saw signatures of such inhibition in the R4d population as well (Figure S5C). R4d population activity in the EB fluctuated in darkness, and similar time-varying activity in darkness was also clearly visible in spontaneous calcium transients in single cells stochastically expressing GCaMP6f (Figure S5H). The time between calcium transients in darkness is relatively regular, lasting between 0 and 3 s (Figure S5I), and centered around 1 s across neurons and across flies (Figure S5J). However, in the presence of a visual stimulus that excites single ring neurons, we saw a consistent reduction in the overall level of R4d ring neuron population activity (Figure S5D,E). The ring neuron that is most strongly excited by the visual stimulus likely inhibits the others, leading to this overall reduction. Such mutual inhibition might produce competitive dynamics between ring neurons (Sun et al., 2017). It may also act as a gain control mechanism that normalizes the net level of visually evoked inhibition produced by the ring neuron population onto the E-PGs in visual panoramas that excite a large population of these neurons (Kim et al., 2019).

Although we did not detect any consistent patterns in the synapse counts between the ring neurons and the E-PGs, we did find an indication of structured inhibition in our calcium activity recordings. In the dark, activity fluctuates between individual ring neurons, suggesting that the whole population is likely active over time, on average. When a stripe is present, however, individual ring neurons appear to suppress the activity of the majority of the ring neuron population. If the weights between individual ring neurons and E-PG neurons are indeed structured to create local troughs of inhibition, signatures of such structure should be visible when an animal transitions from a dark environment (where all of the neurons are active and therefore any circularly permuted structure should average out) to one with a closed loop stripe (where only a small number of ring neurons will be active while the others are suppressed). Indeed, such signatures are visible in the experiments where shi^ts^ was expressed in the E-PG neurons. In these experiments, the E-PG activity localized to one bump at the permissive temperature but was spread throughout the EB at the restrictive temperature (Figure 3F). At the permissive temperature, overall E-PG activity increased when a visual stimulus was present (Figure S4A,E). In contrast, at the restrictive temperature, the overall activity decreased when a visual stimulus was present (Figure S4B,F). These results suggest that the inhibition locally decreases at the position of the bump while globally increasing in the EB, which is consistent with a trough-shaped structure of inhibitory weights between the visual ring neurons and the E-PG neurons.

In summary, R4d ring neuron connectivity to the E-PG neurons, and changes in E-PG activity in response to visual stimulation are both in line with predictions made by ring attractor models of visuomotor integration. Further, unpredicted ring neuron-to-ring neuron connectivity may enable competitive dynamics or gain control across the ring neuron population.

### Pathway from AOTU to EB tethers the compass to visual features

Although we focused on the R4d ring neurons, multiple classes of ring neurons respond to visual features (Omoto et al., 2017). These neurons all arborize in an ancillary structure known as the bulb (BU). They receive visual inputs from a pathway that is conserved across insects and originates in the photoreceptors and travels through the lamina, medulla, and anterior optic tubercle (AOTU) before reaching the BU (Held et al., 2016; Homberg et al., 2011; Homberg et al., 2003). Lineage-based anatomical studies (Omoto et al., 2017) and photoactivatable-GFP (PA-GFP) uncaging in the BU (Sun et al., 2017) suggest that the TuBu neurons provide visual inputs to the ring neurons in the *Drosophila* BU. Silencing the outputs of the TuBu neurons should therefore “blind” the ring neurons and, in turn, “blind” the compass.

Consistent with previous results (Green et al., 2019), in flies with functional AOTU-BU pathways, E-PG neurons maintained a heading representation that accurately tracked a closed-loop stripe, even at high temperatures (Figure 5G). We next expressed shi^ts^ in TuBu neurons to silence their output. At permissive temperatures, flies that expressed shi^ts^ in the TuBu neurons showed normal heading-tuned activity in E-PG neurons during closed-loop walking (Figure S6). At the restrictive temperature, the E-PG population representation was impaired, with an error in tracking heading that matched those observed during walking in darkness (Figure 5H, I bottom, J right). We therefore concluded that the pathway from the AOTU to the BU to the E-PGs (via the TuBu and ring neurons) provides visual cue information that is used as a reference by the fly compass, allowing the fly to maintain arbitrary headings within its surroundings.

### The E-PG, P-EG and P-EN2 excitatory loop that maintains the compass bump in darkness

Finally, we reconstructed two additional classes of columnar neurons that are connected to the E-PG and Δ7 neurons, but whose function in the context of ring attractor models is less obvious: P-EG and N2 neurons (Figure 6A-C) (Green et al., 2017; Wolff et al., 2015). Based on light microscopy and RNA-Seq, P-EG neurons seemed like they could provide direct cholinergic (excitatory) feedback to the E-PG neurons, aiding in the bump’s persistence (Kakaria and de Bivort, 2017). However, although the P-EGs get strong direct input from the E-PGs in the PB (Figure 6D), they do not synapse directly back onto the E-PGs in the EB. Instead they synapse onto the P-EN2s, which are also cholinergic, and in turn synapse onto the E-PGs (Figure 6E) (note that P-EN2s from each side of the PB are densely interconnected with each other and with angular-velocity-carrying P-EN1s in the NO (Figure 6F)). The P-EGs and P-EN2s therefore form a three-synapse feedback loop with the E-PGs: E-PG to P-EG to P-EN2 and back to E-PG. Any P-EG feedback to the E-PGs is therefore mediated by the P-EN2s in the EB. Indeed, P-EN2 activity in the EB largely matches P-EG activity (Figure 6G). Both populations display one bump of activity that moves around the EB, tracking the animal’s heading (Figure 6H), and the bump’s amplitude weakly correlates with the animal’s velocity for each population (Figure S7A-D). The PVA difference between the two is small, around ¼ of the bump full width at half maximum (Figure 6I,J).

**Figure 6:**
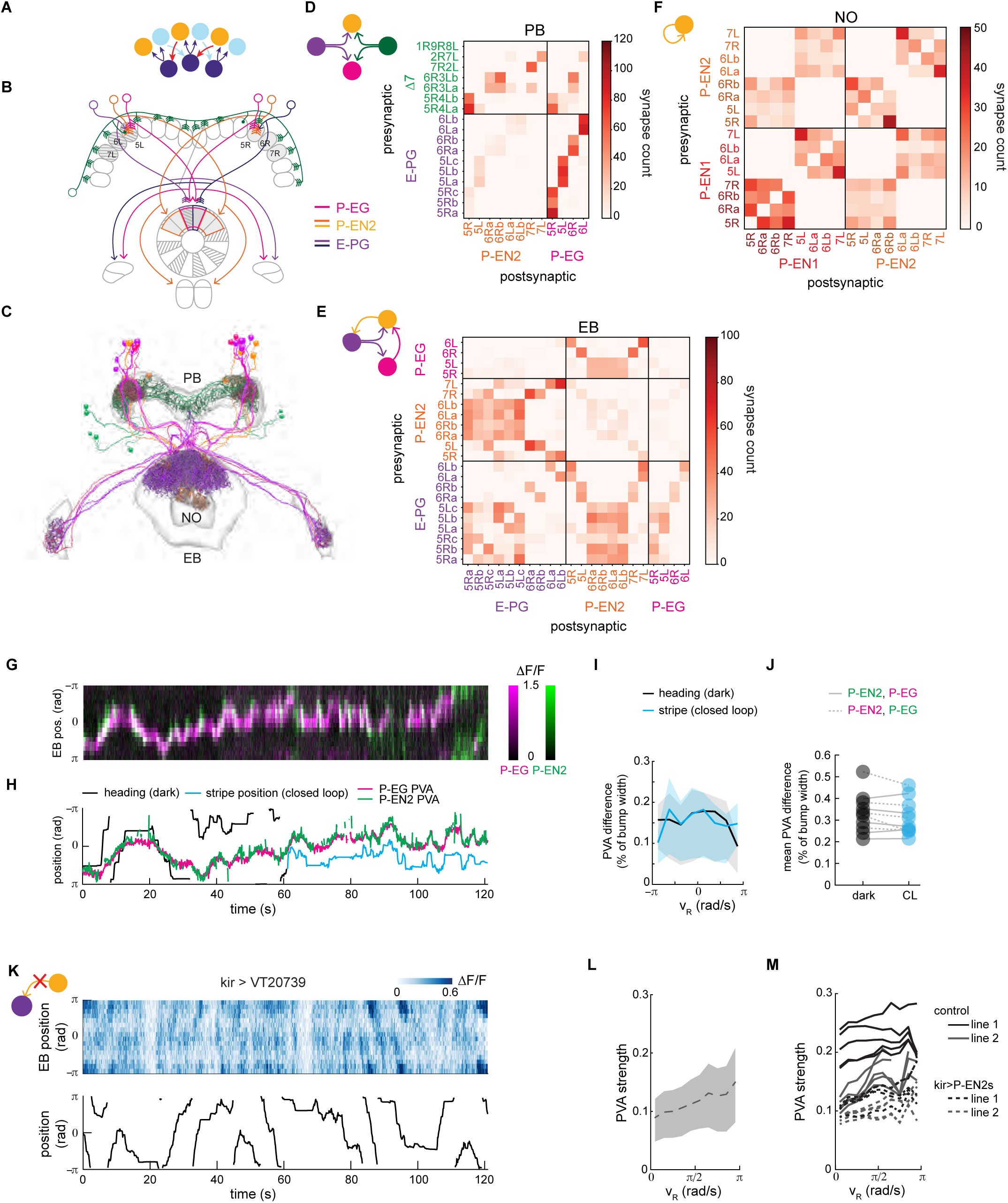
P-EG and P-EN2 neurons together create a distinctive recurrent loop with E-PGs. A. P-EN2 neurons (orange and light blue), like P-EN1s (Figure 4), conjointly encode the fly’s heading and angular velocity, creating an angularly shifted recurrent loop with E-PGs (purple). B. Schematic of P-EG and P-EN2 arborizations in the central brain. EB wedges and PB glomeruli selected for E-PG, P-EG and P-EN2 reconstruction (Figure 6C) are in grey. C. Reconstructed E-PG, P-EG, Δ7 and P-EN2 neurons. Synapses were labelled between these neurons in the NO and Gall and in selected parts of the EB and PB (Figure 6B). D. Connectivity matrix between the E-PGs and Δ7s and the P-EN2s and P-EGs in the PB. The E-PGs form a large number of synapses onto the P-EGs (lower right quadrant). E. Connectivity matrix between the E-PGs, P-EN2s, and P-EGs in the EB. The P-EGs only synapse onto the P-EN2s (top middle) F. Connectivity matrix between the P-EN2s and P-EN1s in the NO. Note that the P-EN1s and P-EN2s are interconnected in each nodulus. G. Two color calcium imaging of P-EG (magenta, labeled with jRGECO1a) and P-EN2 (green, labeled with GCaMP6f) activity in the EB for a fly in the dark. H. The PVAs of P-EG and P-EN2 bumps of activity are shown along with the fly’s heading in the dark (black trace) and the closed-loop stripe position when visible (blue trace). I. Mean PVA difference across rotational velocities for the fly shown in Figure 6F,G. The upper and lower quartile range is shown in the shaded region. J. Mean PVA difference across all velocities across flies in the dark or with a closed-loop stripe. K. (top) Calcium activity in the E-PGs in the EB for a fly in the dark when the P-EN2 neurons express Kir. (bottom) The fly’s heading over the duration of the trial. L. The mean PVA strength of the calcium activity in the EB as a function of the fly’s rotational velocity for the fly shown in H. The standard deviation is shown in the shaded region. M. The PVA strength for 5 flies expressing Kir in the P-EN2s (dotted lines) relative to the control flies (solid lines) as a function of the fly’s rotational velocity.

We next tested the possibility that the P-EN2 population mediates the transmission of delayed excitatory feedback from the P-EGs to the E-PGs, potentially reinforcing the persistence of E-PG heading representation activity. We monitored E-PG activity while silencing P-EN2 activity using Gal80 and the inwardly rectifying potassium channel, Kir (see Methods), thereby breaking the P-EG feedback loop. Consistent with the importance of the P-EG input for maintaining E-PG bump strength, the E-PG bump was almost entirely abolished in the dark in flies in which Kir was expressed in the P-EN2 population (Figure 6K). This was true regardless of the animal’s rotational velocity (Figure 6L,M). Instead, the overall activity in the EB increased as the animal turned (Figure S7J,K). Interestingly, the bump was, however, clearly visible and tracked the fly’s motions in the presence of a stripe in closed loop (Figure S7E-I), consistent with model predictions and recent experiments suggesting that the bump is shaped by visual input from ring neurons (Cope et al., 2017; Fisher et al., 2019; Kim et al., 2019).

In addition to receiving input from the P-EGs in the EB, the P-EN2s also receive input in the PB. There, they receive weak direct E-PG input but strong Δ7 input (Figure 6D). Thus, there is also a second loop to the E-PGs. This second loop goes from E-PGs to Δ7 to P-EN2s and back to E-PGs (Figure 6A). Curiously, this loop anatomically resembles the E-PG and P-EN1 loop, and would not excite E-PG neurons in the same location in the EB. However, a recent study reported that E-PG and P-EN2 activity in the PB are anticorrelated (Green et al., 2017), which we also observed (data not shown). If projected to the EB, this anticorrelated P-EN2 bump in the PB should translate to a bump in the EB that is offset from the P-EG bump, in contrast to what we observed. However, as described above, the P-EN2s also receive direct synaptic input from the P-EGs in the EB, thus complicating the picture in the EB. The discrepancy between P-EN2 activity in the EB and the PB may be accounted for by electrical isolation of the PB and EB compartments of individual P-EN2 neurons (Figure S7N-Q, Supplemental Information). The compartmentalization of P-EN2 activity between the EB and PB raises as-yet-unanswered questions about the functional role of these neurons in the heading direction network.

Similar to the E-PG neurons, the activity of P-EG neurons, which receive considerable synaptic input from E-PGs in the PB, is also offset from the P-EN2 bump in the PB (Figure S7L,M). This offset also matches the difference we see between Δ7 and E-PG activity (Figure 3L). Considering that E-PG neurons provide direct synaptic input to Δ7 neurons, and since those neurons, in turn, make synapses onto P-EN2 neurons, a likely possibility is that the glutamatergic Δ7 neurons excite P-EN2 neurons through excitatory glutamate channels. In contrast, the activity of P-EN1s, whose anatomy is almost identical to the P-EN2s, overlaps with the E-PG activity but is anticorrelated with Δ7 activity. We would therefore expect these neurons to be inhibited by the Δ7s. However, we found no clear difference between glutamate channel mRNA expression in the P-EN1s and P-EN2s (data not shown). Resolving this conundrum will require future experiments that establish the localization of specific glutamate channel types in P-EN1 and P-EN2 to their different arbors in the PB, EB and NO.

In summary, although the P-EN2 offset projection pattern leaves open other possibilities, their role in mediating excitatory feedback from P-EG neurons is likely key to the maintenance of E-PG bump strength in the dark.

### Flies lose flexibility in heading if AOTU input or EB-PB network is disrupted

In our imaging experiments in flies walking in visual VR (previous sections), we focused on how disrupting signals from specific cell types in the heading direction network impacted the stability of the E-PG heading representation. These disruptions also impacted the fly’s behavior. Recent studies have demonstrated that the E-PG compass is required for the fly to select and maintain arbitrary headings relative to visual cues in its surroundings (Giraldo et al., 2018; Green et al., 2019). Consistent with these results, we observed a distinct change in the fly’s fixation preferences when we disrupted the heading representation and its tethering to visual features. (Figure S8A-D; Methods). Control flies displayed individual heading preferences relative to a single stripe that were maintained across trials at both low and high temperatures (Figure S8C left, D, left), but there was a wide range of preferences across flies. A similar range of heading preferences was observed at the permissive temperature in flies that expressed shi^TS^ in the E-PG, TuBu, and Δ7 neurons (Figure S8C top middle, right and Figure S8D top right). In contrast, at the restrictive temperature, we observed these flies all had matching heading preferences, keeping the stripe exclusively in the frontal field of view (Figure S8C bottom middle, right, Figure S8D bottom right). Taken together with previous results (Giraldo et al., 2018; Green et al., 2019), we conclude that flies’ ability to maintain a stable heading representation and exhibit different heading preferences relative to visual cues in their surroundings relies on an intact EB-PB network that receives visual information from the AOTU-BU pathway via the visual ring neurons.

## Discussion

Efforts to model the dynamics of head direction networks have long focused on ring attractors. Although this conceptual framework has been very influential, testing the validity of the structural assumptions and functional predictions made by ring attractor models in biological circuits has been challenging. Here, we applied connectomics, transcriptional profiling, and functional imaging with perturbation of genetically targeted neural populations to examine the structure and function of the neuronal circuits underlying heading direction in the fly. Our results allow us to firmly place several cell types and network motifs from the fly into the context of previous ring attractor models: E-PG neurons construct the heading representation from their inputs; heading-tuned P-EN1 neurons provide angular velocity inputs that update the heading representation when the animal turns; and Δ7 neurons provide inhibition that stabilizes the E-PG heading representation by sculpting P-EN1 activity. We previously invoked the E-PG, P-EN1 and, implicitly, Δ7 neurons in a ring attractor model that captured the observed dynamics of the heading direction network (Figure 7A, see Methods) (Turner-Evans et al., 2017). The weights of this model were tuned without any knowledge of the actual synaptic connections. If we extrapolate the measured synapse counts around the ring to form a complete connectivity matrix of the biological network, the theoretical and the experimentally derived connectivity matrices are similar (Figure 7B). Finally, ring neurons, which are known to bring visual information to the EB in a variety of insects (el Jundi et al., 2014; Heinze and Reppert, 2011; Omoto et al., 2017; Phillips-Portillo, 2012; Seelig and Jayaraman, 2013; Shiozaki and Kazama, 2017; Sun et al., 2017; Vitzthum et al., 2002), make synaptic contacts with E-PG neurons, consistent with recent modeling and experimental work suggesting that the ring neuron population tethers the compass to the fly’s sensory surroundings (Cope et al., 2017; Fisher et al., 2019; Kim et al., 2019).

**Figure 7:**
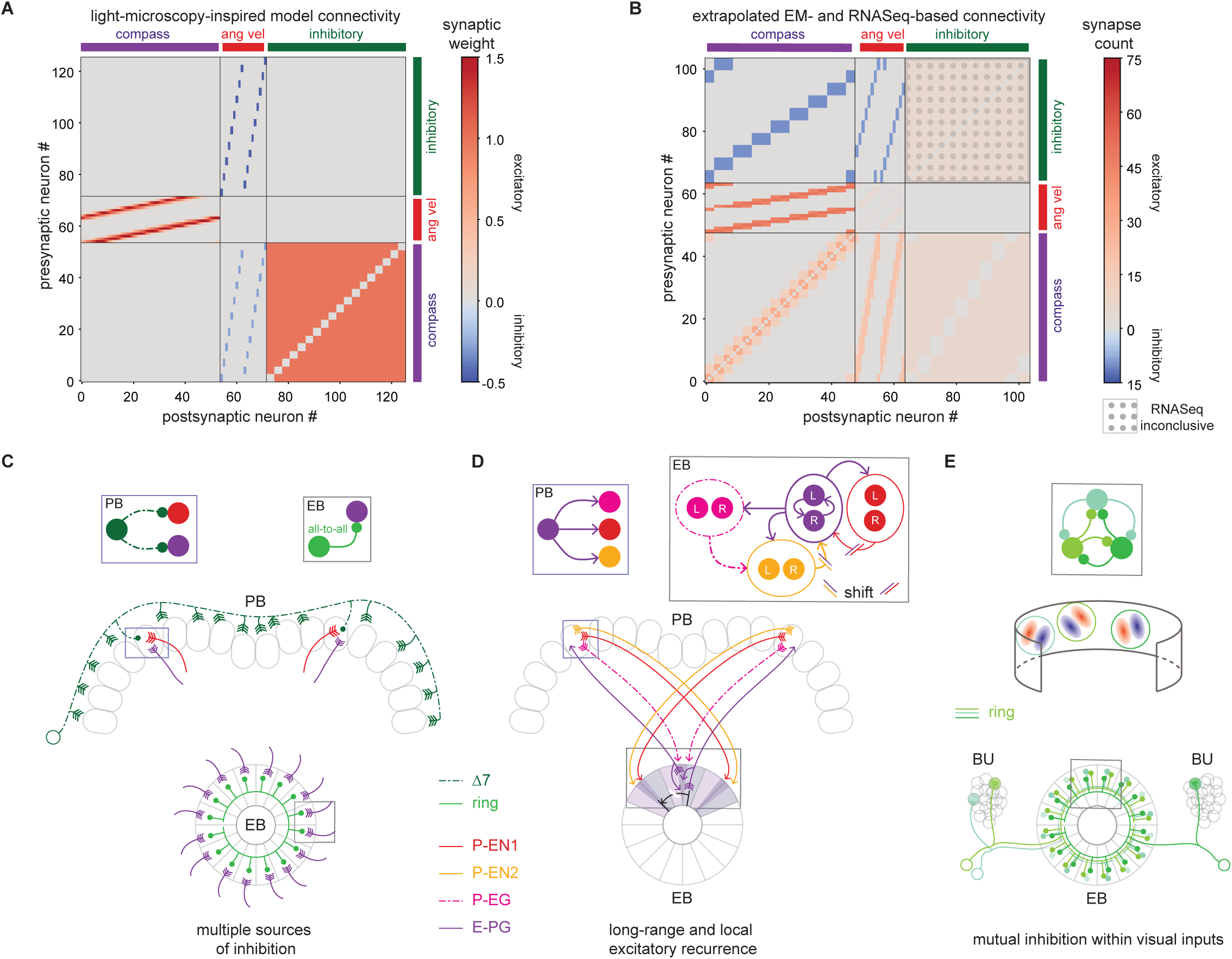
The fly ring attractor network is broadly consistent with theory, but also features unpredicted components and motifs. A. Weight matrix for previous theoretical ring attractor model (Turner-Evans et al., 2017) for the fly compass (see Methods). B. Connectivity matrix extrapolated from data presented in this work. The synapse counts from the EB and PB are averaged and extrapolated to all untraced neurons of every class. Excitatory connections are shown in red and inhibitory connections are shown in blue. C. Inhibition within the fly ring attractor appears to stem from multiple neuron classes, including the Δ7s and ring neurons. The Δ7s provide more structured, long-range inhibition of E-PG and P-EN neurons with different heading tuning in the PB (middle row; expanded view in box at top left), and different classes of ring neurons provide more uniform inhibition of E-PGs in the EB (bottom row; expanded view in box at top right). D. Recurrent excitation between neurons with similar heading tuning (‘local excitation’ in the context of ring attractor models) takes many forms. E-PGs from a specific wedge in the EB synapse onto P-ENs and P-EGs in the PB (schematized in bottom row and expanded into box at upper left). P-ENs project back onto the E-PGs in the EB synapsing onto E-PGs from neighboring tiles (shift shown in bottom row; expanded view in box at upper right). P-EGs also project back without a shift to the EB, where they synapse onto the P-EN2s (upper right). E-PGs also synapse onto P-ENs within the EB (upper right). E. Visual input neurons (simplified receptive fields indicated by red excitatory and blue inhibitory subfields in middle row) synapse onto one another in the EB (bottom row; expanded view in box at top). This mutual inhibition between visually responsive ring neurons may enable winner-take-all dynamics between visual cues, and/or gain control that maintains the same level of inhibition in the circuit in the presence of single or multiple visual cues.

However, while the theoretical predictions and our biological observations are largely consistent, we also uncovered surprises in the biology that will require further theoretical and experimental work to understand. The long-range inhibition that has been invoked to form a stable bump of activity appears to be split across multiple brain regions and multiple classes of neurons (Figure 7C). The Δ7 neurons cannot be the only source of inhibition in the circuit since the E-PG population organizes into a single bump even if the output of the Δ7 neurons is reduced. GABAergic R4d (and other) visual ring neurons also inhibit much of the E-PG population in the presence of visual stimuli. Further, the Gall-EB ring neuron population provides a source of inhibition onto the E-PG neurons (Franconville et al., 2018). We did not reconstruct these neurons, but FISH analysis confirms that they are GABAergic as well (Figure S4L). Indeed, the cumulative inhibition from multiple classes of ring neurons may account for some of the mutual suppression between E-PG neurons (Kim et al., 2017).

Local excitation also appears to be implemented through multiple classes of neurons (Figure 7D). Within the EB, E-PGs synapse onto one another and onto the P-ENs, which in turn synapse back onto the E-PGs—this local recurrence may enable the fly’s internal compass to perform more smoothly than its limited number of neurons may otherwise permit (see, for example, the ‘stickiness’ in bump movement evident in Figure 10 of (Turner-Evans et al., 2017)). Local excitation also appears to be structured across compartments, with the E-PGs synapsing onto the P-ENs in the PB and the P-ENs synapsing on the E-PGs in the EB. Finally, P-EGs appear to create a potentially slower multi-synaptic feedback loop, receiving input from the E-PGs in the PB and synapsing onto the P-EN2s in the EB which in turn synapse back onto the E-PGs. Overall, the redundancy of local excitation and long-range inhibition may allow the network to maintain heading direction activity in the absence of external input. These network features may also provide a means to stabilize the heading representation in the presence of noisy inputs and inhomogeneous synaptic weights, which can disrupt the function of continuous attractor networks (Burak and Fiete, 2012; Compte et al., 2000; Itskov et al., 2011; Renart et al., 2003; Seeholzer et al., 2019; Wu and Amari, 2005; Wu et al., 2008; Zhang, 1996).

Local connections were not limited to the E-PGs. Instead, they appeared across neuron classes and may lead to local computations within the broader ring attractor network. For example, consistent with previous observations from electron microscopy in locusts (Homberg and Muller, 2016) and from *trans-Tango* experiments in flies (Omoto et al., 2018), we observed synapses between the visual ring neurons. These ring neurons tether the bump to a visual scene. Visual scenes are often complex and dynamic and so inter-ring neuron connections may either form a winner-take-all network that leads one visual feature to dominate, preventing the bump from moving erratically as the scene shifts over time (Figure 7E) or provide a mechanism for gain control that normalizes the total level of inhibition from those ring neurons onto the E-PG neurons. The R4d ring neurons traced here are but one of many classes of ring neurons. Other ring neuron types respond to contralateral visual features, or are linked to sleep, circadian rhythm, nutrients, or the animal’s movements (Donlea et al., 2018; Dus et al., 2013; Liang et al., 2019; Liu et al., 2016; Park et al., 2016; Seelig and Jayaraman, 2013; Shiozaki and Kazama, 2017). If similar connections exist between or across these different ring neuron classes, then they may perform computations amongst or between themselves, preprocessing information rather than acting solely as parallel input streams to the E-PG neurons.

Both the structural and functional results also left several puzzles unsolved and raised new questions. The glutamatergic Δ7 neurons synapse onto many columnar neuron types in the PB that project back to the EB, including the P-EN1, P-EN2, and P-EG populations. Indeed, a primary role of the Δ7s may be to pass the heading representation from the E-PGs to other columnar neurons in the circuit. Many columnar neuron types leave the heading direction network and pass into a structure in the CX known as the fan-shaped body. While we did not trace any of these fan-shaped body neurons, we would predict that they also receive input from the Δ7s. Of the P-EN1, P-EN2, and P-EG neurons, it appears that the Δ7 neurons inhibit the P-EN1 and P-EG neurons but may excite the P-EN2 neurons. This excitation would create a bump of P-EN2 activity that is perfectly offset from the E-PG bump in the PB. However, because of compartmentalization, the P-EN2 activity in the PB only produces a small shoulder of a bump in the EB. There, the main part of the P-EN2 activity instead appears to be driven by local excitation from the P-EG neurons at the E-PG bump location. The combination of P-EG and P-EN2 neurons feeding back with a delay onto the E-PG neurons in the EB helps maintain the strength of the heading representation in darkness, but we cannot yet explain the functional role of the shoulder of activity and its displacement from the E-PG bump.

Behaviorally, the heading representation has been shown to be required for the fly to select and maintain arbitrary headings relative to visual cues in its surroundings (Giraldo et al., 2018; Green et al., 2019), a result that we replicated here. Whenever the heading representation was altered, the fly only walked directly towards a visual cue. However, we note that behavioral genetics experiments in the fly have also implicated the CX in sleep and nutrient-state-induced modulation of activity (Donlea et al., 2018; Dus et al., 2013; Liang et al., 2019), short-term visual orientation memory (Kuntz et al., 2017; Neuser et al., 2008), long-term visual and thermal place memory (Liu et al., 2006; Ofstad et al., 2011), and body-size-dependent motor control (Krause et al., 2019; Triphan et al., 2010). The role of the EB-PB circuit in such functions is unknown.

Importantly, the CX and its neurons have been studied not just in other contexts, but also in other insects. Intracellular electrophysiological recordings from CX neurons in a diverse array of insects have discovered topographically-organized visual responses, suggesting the potential involvement of the region in computations ranging from sky-compass-based navigation in locusts (Heinze and Homberg, 2007), monarch butterflies (Heinze and Reppert, 2011), and dung beetles (el Jundi et al., 2015) to optic-flow-based path integration in sweat bees (Stone et al., 2017). Extracellular recording and stimulation in the cockroach CX have also implicated the region in sensory-guided orienting and motor control (Bender et al., 2010; Guo and Ritzmann, 2013; Varga and Ritzmann, 2016). Looking ahead, scaled-up electron microscopic reconstructions of CX networks in different insects (for example, recent EM-guided modeling in the sweat bee CX (Stone et al., 2017)) may enable an exploration of how circuits that are optimized for navigation differ based on the particular sensory statistics, ecological niches and biomechanical constraints of each insect.

It is important to note that the connectivity examined in this study constitutes only a partial connectome, focused on a specific set of neurons selected because of their likely participation in the fly ring attractor network: the E-PG, Δ7, P-EN1, P-EN2, and P-EG neurons. However, this is not an isolated circuit, and we did not attempt to identify all the upstream and downstream partners of these neurons.

Finally, our efforts to link structure to function also highlight the challenges that this task poses. Connectomics is often perceived as providing strong constraints on models of circuit function. However, although this was partly true for the heading direction network, our results also identified unexpected computational mechanisms that the circuit may employ. For example, the profusion of mixed pre- and post-synaptic specializations within single compartments of individual neurons created many more locally recurrent loops between similar and different neuron classes. We cannot yet provide a functional account for such dense recurrence. Future theoretical and experimental work to probe their function may well reveal that the fly brains are more powerful than their numerical simplicity might suggest.

## Methods

### Nomenclature

The columnar neurons (E-PGs, P-EN1s, P-EN2s, and P-EGs) were each assigned a number and an “R” or an “L”. This number-letter combination refers to the glomerulus in the protocerebral bridge in which they arborize. The bridge can be split into right (“R”) and left (“L”) halves, and each half has nine glomeruli (1-9). 9 is always the outer glomerulus while 1 is the glomerulus in the middle. Hence, from left to right, the numbering goes from L9 to L1 and then from R1 to R9. Where multiple neurons of a single type arborize in the same glomerulus, an “a,” “b,” or “c” was appended to the name (e.g. E-PG 5Rc). The Δ7s arborize throughout the bridge but were named based on the 2 or 3 glomeruli to which they send axons. Finally, ring neurons were assigned an arbitrary letter label from a-g.

More information on the anatomy and nomenclature of the individual neurons can be found in (Wolff et al., 2015). Briefly, E-PG and P-EG neurons that arborize in glomerulus 5 (right or left) project to the 12 o’clock position in the ellipsoid body. P-EN1s and P-EN2s project to offset positions in the ellipsoid body (10:30 for the right bridge neurons, and 1:30 for the left bridge neurons). This offset is consistent for all similar E-PG/P-EG and P-EN1/P-EN2 pairs.

### Experimental protocols

#### Genotypes

| **Experiment** | **Parent #1** | **Parent #2** |
| --- | --- | --- |
| **FISH** | UAS-HaloTag | SS00096 |
| " | " | SS00131 |
| " | " | GR37F06 |
| " | " | VT08135 |
| " | " | SS02238 |
| " | " | SS30295 |
| " | " | SS02232 |
| " | " | SS02233 |
| " | " | SS02191 |
| " | " | SS27853 |
| " | " | Ring #1 |
| " | " | Ring #2 |
| **FLP-out GCaMP** | 20XUAS>myrTopHat2>Syn21-OpGCaMP6f-p10 su(Hw) attP8; pGP-JFRC59-13XLexAop2-IVS-NES-jRGECO1a-p10 in su(Hw)attP5; + | pBhsFlp2::PEST; LexA-VT25957;EB1 |
| " | " | pBhsFlp2::PEST; LexA-VT25957;c232 |
| **Two color calcium imaging** | pGP-JFRC59-13XLexAop2-IVS-p10-GCaMP6f; 20XUAS-IVS-jRGECO1a-p10 | LexA-60D05; GR55G08 |
|  | " | LexA-55G08; R60D05 |
|  | 13XLexAop2-IVS-Syn21-OpGCaMP6f-p10 su(Hw) attp40; LexA-37F06 VK22; 20XUAS-IVS-jRGECO1a-10 | SS02191 |
|  | 20XUAS-IVS-Syn21-OpGCaMP6f p10 su(Hw)attP8; LexA-37F06 VK22; 13XLexAop2-IVS-Syn21-jRGECO1a | SS02191 |
| **Shi** | LexA-VT25957; pJFRC100-20XUAS-TTS-Shibire-ts1-p10 in VK00005, 13xLexA-syn21-syt-OpGCaMP6f-p10 in attP2 (wtB) | SS00096 |
|  | " | LexA-60D05; GR55G08 |
|  | " | R76B06 |
| **Kir** | tubP Gal80ts; UAS-kir2.1::GFP | LexA-27F02, LexAOp-jRGECO1a; VT20739 |
|  | " | LexA-27F02, LexAOp-jRGECO1a; GR12D09 |
| **RNA-Seq** | 10XUAS-IVS-myr::GFP; 10XUAS-IVS-nls-tdTomato | SS000096 |
|  | " | SS000131 |
|  | " | SS30295 |
|  | " | SS02238 |
|  | " | SS02191 |
|  | " | SS00238 |
|  | " | SS02232 |
|  | " | SS62862 |

#### Simultaneous behavioral and calcium activity recordings of a fly on a ball

All imaging experiments were performed with head-fixed, tethered female flies walking on an air-suspended ball as described in (Seelig et al., 2010). Each fly was first cold anaesthetized and the proboscis fixed with wax to limit brain movement. The fly was then glued to a pin and positioned in an inverted pyramidal metal shim such that the top of the head was exposed while the body was free and visibility only minimally occluded. The head was then glued to the shim and immersed in saline. The cuticle was then removed above the brain, along with any fat, and the air sacs and trachea were pinned to the side. Finally, the muscles that run along the top of the esophagus were cut to limit brain movement. The holder was then positioned such that the fly was centered on the ball and could walk freely.

The movements of the ball (a proxy for the walking movements of the fly) were recorded using the fly spherical treadmill system (Seelig et al., 2010). These movements were fed via USB to a PC and both recorded and used to update closed loop visual stimuli as needed (described further below).

Calcium imaging experiments were performed as described in (Turner-Evans et al., 2017). Briefly, a custom two-photon microscope running ScanImage, similar to the one described in (Seelig and Jayaraman, 2015), was used for all experiments. A full description of the filter sets can be found in (Turner-Evans et al., 2017). All imaging was performed using a tunable femtosecond pulsed laser at ∼20 mW power, set at 930 nm for one color and 1020 for two color experiments. 256 × 256 pixel images were obtained across multiple Z planes to form a volume. The volumes were ~50 μm on a side and consisted of 6 Z planes, each spaced 8 μm apart (for ellipsoid body imaging) or 12 Z planes, each spaced 4 μm apart (for protocerebral bridge imaging). Volumes were obtained at a rate of 11.4 Hz and 8.2 Hz, respectively. The ball position was tracked at 200 Hz and synchronized to the calcium imaging using TTL pulses triggered after each frame grab.

For the single ring neuron FLP-out experiments, high resolution stacks were taken at the end of every experimental session to inspect the ring anatomy. These stacks were 1024 x 1024 pixels, with 25 planes each spaced 2 μm apart, and the laser power was turned up to ∼100 mW to obtain 20 volumes with these parameters.

#### Open and closed-loop visual display

A custom-built digital micromirror device (DMD) projector system was used to show visual stimuli to the flies. A 240°, 2.5 inch tall, 3 inch diameter, cylindrical frame was 3D printed and covered with a single white diffuser sheet (V-HHDE-PM06-S01-D01 sample, BrightView Technologies, Durham, NC, USA). Three projectors each covered 80° of the screen (two DepthQ WXGA 360 HD 3D Projector, developed by Anthony Leonardo, Janelia, and Lightspeed Design, Bellevue, WA, USA and one Texas Instruments DLP 3010 Light Control). These projectors were modified to accommodate external blue LEDs as their light source (ThorLabs M455L3 455 nm LEDs with two Semrock BrightLine 447/60 nm bandpass filters in the path)

Custom C++ code using the OpenGL API generated the visual projections, which was either a completely dark virtual arena or a 15° bright stripe on a dark background. The projections were rendered with an Nvidia Quaddro K4000 graphics card. The code creates virtual objects and then uses the treadmill positional data to update the objects’ positions as the fly moves. Brightness and spatial distortions due to the cylindrical screen are corrected via the software.

The voltage from a photodiode placed on top of the screen was recorded with a DAQ each time the projection was updated (at a 360 Hz framerate). The DAQ also recorded a TTL pulse each time a calcium imaging frame was grabbed. The two signals were then registered with respect to one another to link activity and behavior.

Visual stimuli consisted of one of four different conditions: darkness, a closed-loop stripe, an open loop stripe that circled the projection screen every 5 s, or a closed-loop stripe that periodically underwent open-loop jumps of 120°. For the closed-loop stripe trials, whenever the stripe moved into the 120° arc behind the fly where no screen or projector was present, it was moved such that it immediately appeared on the other side, making the closed-loop span effectively 240°.

#### Shi experiments

Experiments using the temperature sensitive, synaptic vesicle reuptake inhibitor Shibire^TS^ were performed as described in (Turner-Evans et al., 2017). Briefly, the first two trials occurred at room temperature (the permissive temperature). Heated saline was then perfused across the brain until a temperature of 30°C (the restrictive temperature) was reached, as measured by a thermocouple placed near the head capsule. The fly was then shown a closed-loop 15° stripe for 10 min, during which time the temperature was stabilized at 30°C. Three trials were then performed at the elevated temperature.

#### Kir experiments

At low temperatures, Gal80ts inhibits Gal4 transcriptional activity. At high temperatures, Gal80ts becomes inactive, leading to kir transcription. Flies were reared and raised for one day at 21.5°C before being split into two groups. Group one (the control group) was maintained at 21.5°C for two more days while group two (the experimental group) was shifted to a 30°C incubator. At 30°C, Gal80ts becomes inactive, allowing kir to be transcribed. Immediately before each experiment, either control or experimental flies were taken out of the incubator and dissected and measured at room temperature.

#### Electron microscopy reconstruction: Neuron Tracing

Neuron reconstructions were created from manual skeleton tracing of neuronal arbors and annotation of synapses from image stacks in CATMAID (http://www.catmaid.org) as described in Zheng et al. (2018). In brief, all neurons included in analyses were reconstructed by a minimum of two team members, an initial tracer and a subsequent proofreader who corroborated the tracer’s work. To parallelize reconstruction efforts, some neurons were assigned more than one initial tracer and/or more than one proofreader. If a tracer or proofreader encountered ambiguous features (neural processes or synapses that were not identifiable with high confidence), (s)he consulted other tracers and proofreaders to determine the validity of the features in question, enlisting more expert tracers as needed. Any feature that remained ambiguous after scrutiny by an expert tracer was not included in the neural reconstruction and/or flagged to be excluded from analyses. During the proofreading phase, the proofreaders and tracers iteratively consulted each other until each neuron was deemed morphologically and synaptically correct per its tracing objective. Chemical synapse identification required that at least three of the following features were present, with the first being required: 1) an active zone including vesicles; 2) presynaptic specializations (i.e. a ribbon or T-bar with or without a platform); 3) synaptic clefts; 4) postsynaptic membrane specializations such as postsynaptic densities (PSDs). In cases where PSDs were not visible, all cells with membranes that had unobstructed access to a clearly visible synaptic cleft were annotated as postsynaptic. Since gap junctions are unlikely to be resolved at the resolution of this dataset we made no effort to annotate them.

#### Electron microscopy reconstruction: Tracing to classification

Neural reconstruction of microtubule-containing backbone neurites (see Schneider-Mizell et al., 2016) or gross morphology alone is often sufficient to classify a neuron based on expert identification or searching of existing LM datasets with NBLAST software (Costa et al., 2016). Neurons traced to classification were at a minimum reconstructed, with or without synapses, to the point at which their gross morphologies (i.e. backbone skeletons) unambiguously matched that observed by LM for a given cell class, or were unambiguously deemed to be a new cell type not previously observed.

#### Electron microscopy reconstruction: Tracing to completion

Every identifiable neurite was traced to achieve morphological completion, and every identifiable presynaptic release site or postsynaptic density was annotated to achieve pre- or postsynaptic completion, respectively, for a given neuron. Some neurons were traced to morphological or synaptic completion only on portions of their arbors or only in certain brain regions (e.g., D7 presynaptic glomeruli were traced to morphological and synaptic completion, whereas the remainder of these neurons were only traced to classification).

#### RNA Sequencing: Expression checks

Neurons of interest were isolated by expressing the fluorescent reporter 10XUAS-IVS-myr::GFP; 10XUAS-IVS-nls-tdTomato using split-Gal4 drivers specific for particular cell types and then manually picking the fluorescent neurons from dissociated brain tissue. As a preliminary to the sorting process, each driver/reporter combination was ‘expression checked’ to determine if the marked cells were sufficiently bright to be sorted effectively and if there was any off-target expression in neurons other than those of interest. Drivers that met both these requirements were used in sorting experiments as described below.

#### RNA Sequencing: Sorting of fluorescent-labelled neurons

*Drosophila* adults were collected daily as they eclosed, and aged 3-5 days prior to dissection. For each sample, 60-100 brains were dissected in freshly prepared, ice cold Adult Hemolymph Solution (AHS, 108 mM NaCl, 5 mM KCl, 2 mM CaCl_2_, 8.2 mM MgCl_2_, 4 mM NaHCO_3_, 1 mM NaH_2_PO_4_, 5 mM HEPES, 6 mM Trehalose, 10 mM Sucrose), and the major tracheal branches removed. The brains were transferred to a 1.5 ml Eppendorf tube containing 500 microliters 1 mg/ml Liberase DH (Roche, prepared according to the manufacturer’s recommendation) in AHS, and digested for 1 h at room temperature. The Liberase solution was removed and the brains washed twice with 800 microliters ice cold AHS. The final wash was removed completely and 400 microliters of AHS+2% Fetal Bovine Serum (FBS, Sigma) were added. The brain samples were gently triturated with a series of fire-polished, FBS-coated Pasteur pipettes of descending pore sizes until the tissue was homogenized, after which the tube was allowed to stand for 2-3 m so that the larger debris could settle.

For hand sorting, the cell suspension was transferred to a Sylgard-lined Glass Bottom Dish (Willco Wells), leaving the debris at the bottom of the Eppendorf tube, and distributed evenly in a rectangular area in the center of the plate with the pipet tip. The cells were allowed to settle for 10-30 m prior to picking. Fluorescent cells were picked with a mouth aspirator consisting of a 0.8 mm Nalgene Syringe Filter (Thermo), a short stretch of tubing, a plastic needle holder, and a pulled Kwik-Fil Borosilicate Glass capillary (Fisher). Cells picked off the primary plate were transferred to a Sylgard-lined 35 mm Mat Tek Glass Bottom Microwell Dishes (Mat Tek) filled with 170 microliters AHS+2%FBS, allowed to settle, and then re-picked. Three washes were performed in this way before the purified cells were picked and transferred into 50 microliters buffer XB from the PicoPure RNA Isolation Kit (Life Technologies), lysed for 5 m at 42°C, and stored at −80°C.

For FACS, the samples were triturated in AHS+2%FBS that was run through a 0.2 micron filter, and the cell suspension was passed through a Falcon 5 ml round-bottom tube fitted with a 35 micrometer cell strainer cap (Fisher). Samples were sorted on a SONY SH800 cell sorter gated for single cells with a fluorescence intensity exceeding that of a non-fluorescent control. For bulk samples, GFP+tdTom+ events were purity sorted directly into 50 microliters PicoPure XB buffer, the sample lysed for 5 m at 42°C, and stored at −80°C. For low-cell samples, replicates of 40 cells were either purity sorted directly or yield sorted to enrich for GFP+tdTom+ cells followed by a purity sort into 3 μL Lysis Buffer (0.2% Triton X-100, 0.1 U/μL RNAsin) and flash frozen on dry ice. For single-cell (scRNA-Seq), individual cells were purity sorted directly into a 96-well plate containing 3 μL Lysis Buffer per well.

#### RNA Sequencing: Library preparation and sequencing

For bulk samples, total RNA was extracted from 100-500 pooled cells using the PicoPure kit (Life Technologies) according to the manufacturer’s recommendation, including the on-column DNAse1 step. The extracted RNA was converted to cDNA and amplified with the Ovation RNA-Seq System V2 (NuGEN), and the yield quantified by NanoDrop (Thermo). The cDNA was fragmented and the sequencing adaptors ligated onto the fragments using the Ovation Rapid Library System (NuGEN). Library quality and concentration was determined with the Kapa Illumina Library Quantification Kit (KK4854, Kapa Biosystems), and the libraries were pooled and sequenced on an Illumina NextSeq 550 with 75 base pair reads. Sequencing adapters were trimmed from the reads with Cutadapt (Martin, 2011) prior to alignment with STAR (Dobin et al., 2013) to the *Drosophila* r6.17 genome assembly (Flybase). The resulting transcript alignments were passed to RSEM (Li and Dewey, 2011) to generate gene expression counts. For low-cell and single-cell samples, one microliter of harsh lysis buffer (50 mM Tris pH 8.0, 5 mM EDTA pH 8.0, 10 mM DTT, 1% Tween-20, 1% Triton X-100, 0.1 g/L Proteinase K (Roche), 2.5 mM dNTPs (Takara), and ERCC Mix 1 (Thermo-Fisher) diluted to 1e-7) and 1 microliter 10 μM barcoded RT primer were added, and the samples were incubated for 5 m at 50°C to lyse the cells, followed by 20 m at 80°C to inactivate the Proteinase K. Reverse transcription master mix (2μL 5X RT Buffer (Thermo-Fisher), 2 μL 5M Betaine (Sigma-Aldrich), 0.2 μL 50 μM E5V6NEXT template-switch oligo (Integrated DNA Technologies), 0.1 μL 200 U/μL Maxima H-RT (Thermo-Fisher), 0.1 μL 40U/μL RNAsin (Lucigen), and 0.6 μL nuclease-free water (Thermo-Fisher) was added to the lysis reaction and incubated at 42°C for 1.5 h, followed by 10 m at 75°C to inactivate the reverse transcriptase. PCR was performed by adding 10 μL 2X HiFi PCR Mix (Kapa Biosystems) and 0.5 μL 60 μM SINGV6 primer and incubating at 98°C for 3 m, followed by 20 cycles of 98°C for 20 s, 64°C for 15 s, 72°C for 4 m, and a final extension of 5 m at 72°C. Groups of 8 reactions were pooled to yield ∼250 μL and purified with Ampure XP Beads (0.6x ratio; Beckman Coulter), washed twice with 75% Ethanol, and eluted in 40 μL nuclease-free water. The DNA concentration of each sample was determined using Qubit High-Sensitivity DNA kit (Thermo-Fisher).

To prepare the Illumina sequencing library, 600 pg cDNA from each pooled sample was used in a modified Nextera XT library preparation (Illumina) (Soumillon et al., 2014) using the P5NEXTPT5 primer and extending the tagmentation time to 15 m. The resulting libraries were purified according to the Nextera XT protocol (0.6x ratio), and quantified by qPCR using Kapa Library Quantification (Kapa Biosystems). Six to eight sequencing libraries were loaded on a NextSeq High Output flow cell reading 25 bases in read 1, including the spacer, sample barcode and UMI, 8 bases in the i7 index read, and 75 bases in read 2 representing the cDNA fragment from the 3’ end of the transcript.

#### Fluoresence *in Situ* Hybridization

Fluorescence in situ hybridization followed the protocol outlined in (Meissner et al., 2019)

#### Stochastic expression of GCaMP6f in ring neurons with the FLP-out technique

Flies were reared at 25°C. Less than one day after eclosion, adults were heat shocked in a water bath at 37°C for 8 min, which consistently lead to 0-3 individual neurons to be labeled with GCaMP.

### Simulation

#### Biophysical modelling of P-EN2s

All biophysical modelling was done using the NEURON with python software (Hines et al., 2009). For the simulations, neurons of interest were first imported into python using pymaid (https://pymaid.readthedocs.io). They were then simplified by removing all stretches of nodes that were not branchpoints or synapses and replacing them with a single straight cylinder of the same length. The width of the cylinder was taken to be *d* = 0.442 *μm* for axonic regions and *d* = (0.442 − 0.025)e^−0.040589l^ + 0.025 for dendritic regions, where d is given in *μm* and *l* is measured in nm. These values were based on empirical data from the EM images. For all simulations, we set *E_rest_* = −55 *mV*, R_m_ = 20800 ΩCm^2^, C_m_ = 0.79 uF cm^-2^, and R_a_ = 266Ωcm (Gouwens and Wilson, 2009). We implemented all synapses as NEURON Exp2Syn synapses and let cholinergic synapses have a reversal potential of *E_rev_* = 0 mV, time, constants of *τ*_1_ = 0.61 *ms* and *τ*_2_ = 2.11 *ms*, and a maximum conductance of 16/75000 *μS* constants of (Lee and O’Dowd, 1999).

To quantify the variability in potential within and between neuropils in P-EN2 neurons for a given number of input synapses *n*, we first selected *n* Δ7 input synapses at random 10 times in each of the 5 P-EN2 neurons for which we have traced Δ7 input neurons. We then ran a NEURON simulation for 70ms with each input synapse firing thrice at 25ms intervals with a NetStim noise parameter of 0.2. In the EB, we quantified the potential at all E-PG output synapses and found the maximum potential for each synapse. We then averaged the mean and standard deviation of these maximum potentials across all simulations for a given *n*. In the PB we averaged the maximum potentials over all input synapses from Δ7 neurons since the P-EN2 neurons do not have significant output in the PB.

#### Extrapolation of the ring attractor weight matrix from Turner-Evans*, Wegener*, et al.

The rate model in (Turner-Evans et al., 2017) only explicitly included the E-PGs and P-EN1s. Inhibition was assumed to come from the Δ7s, but it was incorporated into the E-PG connectivity matrix. In this matrix, each E-PG inhibited all P-EN1s. However, the inhibition weights were not uniform. One P-EN1 was less strongly inhibited than the rest. The location of the less inhibited P-EN1 was chosen such that it matched up with a particular E-PG. Both that E-PG and its partner P-EN1 were assumed to code for the same heading direction angle.

For Figure 7A, the Δ7s were instead assumed to act as a standalone population. In this scenario, each Δ7 inhibits one P-EN1 and receives excitatory input from all E-PGs expect for those that code for the same heading direction angle as the chosen P-EN1. In the limit that the Δ7 time constant goes to 0, the weight matrix in Figure 7A will match that of (Turner-Evans et al., 2017).

### Data Analysis

#### Calcium imaging and behavior preprocessing

Maximum intensity projections were obtained for each volume within a stack. To correct for movement, a mean of the individual projections was then computed, and each projection was registered to the mean through normalized cross correlation. In the case of two-color imaging, the green channel was used for registration.

Reference regions of interest were defined for each fly by drawing polygons over the mean maximum intensity projection of a chosen stack. For two-color imaging a composite of the two channels was used. For protocerebral bridge imaging, eighteen polygons were placed over the structure, one for each glomerulus. For ellipsoid body imaging, ellipses were drawn over the inner and outer circumference of the structure and the inner region equi-angularly divided into 16 regions. The reference regions of interest were then individually placed over each mean projection of each stack for each experiment.

The registered stacks were then Gaussian filtered with a 2×2 pixel size and a standard deviation of 0.5 pixels. The normalized change in fluorescence for each region was then calculated as follows. The baseline fluorescence was taken as the mean of the lowest 10% fluorescence values over the experiment. All fluorescence values were then normalized by this value and then 1 was subtracted from them.

The recorded behavioral data was then registered to the imaging stacks by way of the frame grab and photodiode signals (see third paragraph of open and closed-loop visual display above).

#### Population vector average (PVA)

The MATLAB-based Circular Statistics Toolbox (Berens, 2009) was used to calculate the population vector average (PVA) and the population vector average strength. The PVA is the circular mean (circ_mean) of the 16 ROIs while the PVA strength is their resultant vector length (circ_r).

#### Unwrapping heading and PVA signals

The heading and PVA signals were unwrapped by finding time points where the signal jumped by more than π or 2π/3, respectively. The signal was then shifted up or down by 2π at those /3 points, as appropriate.

#### Quantifying bump stability

We quantified the bump’s stability as follows. First, we plotted how much the bump’s angular orientation changed versus how much the fly’s heading direction changed at every time point in a 4 s sliding window (see Figure S1C-E). To pick the time window for this analysis, we first plotted the median correlation between bump and fly orientation within different time windows. We picked the time window for which the correlation values began to saturate for all the conditions that we analyzed (darkness and closed-loop; permissive and restrictive temperatures in control and Shi flies), but during which time the bump could still be tracked (for some Shi experiments, the bump became difficult to track for longer intervals at the restrictive temperature). We then fit a line to these points and calculated the slope. The slope was the gain between the fly’s turns and the bump’s movements. A gain of 1 implied that the bump perfectly tracked the fly’s heading direction.

#### AOTU Shi knockout analysis: Calcium activity and behavior

The offset between the bump position and the fly’s heading was calculated by taking the difference between the PVA and the heading. This offset was then binned into 17 linearly spaced bins between −π and π. The mean resultant vector length was then calculated using the circ_r function described above. The behavior was binned and analyzed in the same way, though now the stripe position was used.

#### Electron microscopy skeleton rendering

Traced neurons were imported into Blender using Philipp Schlegel’s CATMAID-to-Blender plugin (https://github.com/schlegelp/CATMAID-to-Blender) (Schlegel et al., 2016). Classes were then separated into layers; their synaptic inputs and outputs were removed; and they were given unique colors. The bevel depth (spline thickness) was increased and shaded using a Strahler analysis (alpha decreases as branch order increases away from root node). Finally, cameras were snapped to the neuron bounding dimensions, a hemi lamp was added, and the resulting images were rendered as 200% resolution png files.

#### Connectivity matrices and distributions

Analyses were carried out in python after importing the neurons using pymaid. For all neuropil-specific analyses, neurons were divided into distinct regions at points along the axon approximately halfway between each pair of neuropils. Connectivity matrices were generated in python using pyplot’s imshow function and the ‘Reds’ colormap.

In the following, we use uppercase letters (*A, B*) to refer to types of neurons, and lowercase letters (*a, b*) to refer to single neurons of type *A/B*. The fraction of input synapses to a neuron *a* that come from a given type of neurons *B* was calculated as the number of annotated synapses from *B* to *a* divided by the total number of annotated postsynaptic sites on *a*. The fraction of output synapses from *a* to *B* was calculated similarly.

#### Estimates of total connectivity distributions

Estimates of the total proportion of connectivity accounted for by the cell types considered in the present manuscript were based on light-level anatomy from Wolff et al. (2015). We assume that the central complex contains 16 P-EN1, P-EN2, and P-EG neurons, 54 E-PG neurons, 40 Δ7 neurons, and 100 ring neurons. In the noduli, we assume that there is no compartmentalization and that neurons are distributed evenly between the left and right noduli. We treat the left and right noduli separately. We thus take the estimated connectivity from a neuron *a* to a neuron *b* in a nodulus to be the mean fraction of output synapses from a neuron *a’* that go to a neuron *b’*, averaged over all pairs of *A* and *B* neurons, times the total number of presynaptic sites in neuron *a*. To estimate the number of post-synapses in *b* that come from *a,* we take the mean number of synapses in *b’* that come from *a’*, averaged over all pairs, and multiply by the total input to *b*. For the total output from neuron *a* to any neuron of type *B*, we multiply the connectivity from neuron *a* to neuron *b* by the total number of neurons of type *B* in a nodulus. For the compartmentalized neuropils discussed below, we instead multiply the compartment-specific average connectivity by the total number of neurons of the relevant type in the compartment.

In the PB, we assume that connections between P-EN, E-PG and P-EG neurons only occur within a single glomerulus. We take the average connectivity between each pair of these types of neurons to be the average over all glomeruli for which we have data, assuming 1 P-EN/P-EG neuron, 3 E-PG neurons, and 5 Δ7 neurons per glomerulus when calculating total connectivity. We assume that each Δ7 neuron gives output to two glomeruli where it does not receive any input and that the Δ7 input is uniformly distributed across the remaining glomeruli. This violates the known anatomy of the Δ7 neurons but provides a good approximation to average and total connectivities given the limited number of glomeruli from which we have data. When estimating Δ7-Δ7 connectivity, we do not consider any compartmentalization.

In the EB, we estimate connectivities to and from E-PG neurons at the level of wedges and connectivities between P-EG and P-EN neurons at the level of tiles. Each tile covers 45° of the EB. For tile neurons, we consider the total connectivity to be the sum of the total connectivity within the same tile and the connectivity to and from the two neighboring tiles. For E-PG neurons, we similarly consider connectivity within the same wedge and to the two neighboring wedges in the case of connectivity with wedge neurons, and tiles in the case of connectivity with tile neurons. For a given pair of neuron types and compartment or pair of compartments, connectivity is estimated as described for the noduli above. We assume ring neurons to be uniformly distributed throughout the EB and do not distinguish between different classes of ring neurons.

#### RNA Seq analysis, plotting

Counts were normalized to the number of genic reads per million reads.

#### E-PGs: Shi experiments

The PVA strength was calculated as stated above, and the mean of the PVA strength was then found across the entire trial. For Figure S4G, the stripe position was distributed between 60° bins. The mean and standard deviation of the activity were then calculated across all time points within these bins and plotted.

#### Δ7s and E-PGs: Two color calcium imaging analysis

For the registered bump activity plots shown in Figure 3K, the maximum of the red channel was found at each time point, and the activity profiles of the red and green time channels across ROIs were each circularly shifted by the same number of glomeruli so that red channel peak was positioned at glomerulus 4. These profiles were then sorted by rotational velocity, across 7 bins between −180 and 180°/s. The mean PVA was then found for these sorted profiles, and the difference between them calculated and plotted in Figure 3L.

#### Δ7s and E-PGs: Shi experiments

A 4 s sliding window was moved over the PVA and heading data. If the PVA strength was over 0.075 within that window (meaning that a clear bump of activity was present), the PVA was plotted with respect to the heading, and the slope and correlation coefficient were calculated. These slopes were then placed into one of 21 linearly spaced bins between −4 and 4. The median slope and the 25^th^ to 75^th^ percentile range were then calculated.

#### Fluorescence *in Situ* Hybridization data analysis

63x images were manually inspected for overlap between the GFP tagged cell bodies and the tagged neurotransmitter specific mRNA.

#### FLP-out ring neuron analysis

Five trials were run for each fly. The activity was then averaged over these trials. Open-loop responses were further averaged over all six clockwise or counterclockwise repetitions. The maximum or minimum activity was then found for these averaged signals.

Dark activity of single ring neurons was deconvolved into events by using the fast oopsi non-negative deconvolution algorithm (https://github.com/jovo/fast-oopsi) (Vogelstein et al., 2010). The time between events was then binned into one of 25 linearly spaced bins in between 0 and 3 s.

#### Population ring neuron analysis

Regions of interest were manually drawn over the center ring and on between 2 and 4 visible glomeruli on each side in the BU. The signal in these ROIs was then averaged over the 5 trials.

#### P-EN2s and P-EGs: Two-color calcium imaging analysis

The PVA was calculated for both the P-EN2s and the P-EGs. Time points were then binned by rotational velocity (in one of eight equally spaced bins between -π and πrad/s). At times when the PVA strength was greater than 0.1, the difference between the P-EG and the P-EN2 PVA was calculated and the mean and standard deviation calculated over all points within a given bin. The full-width-at-half-maximum (FWHM) of the activity was similarly calculated at each point in time and binned by rotational velocity. The mean FWHM for each bin was then calculated. The mean and SD of the PVA difference in each bin was then divided by the corresponding mean FWHM to arrive at the plot in Figure 6H. For the plot in Figure 6I, each individual PVA difference was divided by the FWHM that fell within its rotational velocity bin and all of these width normalized values were then averaged.

Cross-correlation between the maximum red or green ΔF/F and the rotational or forward velocity was calculated using the corrcoef function in MATL for each time lag (Figure S7A-D).

#### P-EN2s and P-EGs: Kir experiments

The PVA strength of the signal was calculated at each time point and binned according to the rotational speed (in one of 11 equally space bins between 0 and π). The mean and standard deviation were then calculated over all points within a given bin.

The mean ΔF/F for the off-peak ROIs was calculated as follows: the maximum ΔF/F ROI was found, and that ROI and one on either side of it were excluded from the analysis. The mean of the remaining 13 ROIs was then found and binned and analyzed as above for the PVA Strength (Figure S7J,K)

#### P-EN2 and E-PG: EB projections

The signals were registered to the red, E-PG peak and sorted by rotational velocity as outlined in the “Δ7s and E-PGs: Two color calcium imaging analysis” section.

## Acknowledgments

We thank Ruchi Parekh and many members of Janelia’s CAT for help with reconstructing neurons, in particular: Marisa Dreher, Chris Patrick, Tansy Yang, Cullen Moran, John Walsh, Neha, Kelli Fairbanks, Mia Ndama and Maggie Greer. We are also grateful to interns, Claire Peterson, Stanley Tran and Boyd Qu for neuron tracing. We thank the Janelia Quantitative Genomics team for performing RNA sequencing and Vilas Menon for helpful advice on RNA-Seq data analysis, and Geoffrey Meissner for carrying out the FISH experiments. We are grateful to Romain Franconville for his help in selecting Gal4 lines suitable for FISH and RNA-Seq, to Ed Rogers, Karen Hibbard, and Michael Rogers for generously providing fly constructs, and also for advice on fly wrangling. We thank Scarlett Harrison, Guillermo Gonzolez, Brenda Paez and Todd Laverty for help with fly husbandry; Vasily Goncharov, Chris McRaven, and Jon Arnold for assistance with the two-photon microscope; Bill Biddle for hardware assistance for the VR setup; and Jim Strother for initial code for the VR setup. We thank Brad Hulse for initial code and discussion on bump stability analyses; Hannah Haberkern for insights on behavior. Finally, we thank Sung Soo Kim, Chuntao Dan, Ann Hermundstad, Romain Franconville, Brad Hulse, and Hannah Haberkern for all around good advice and feedback!

## Supplemental Information

### Ring neurons may also communicate with the E-PGs via wake-promoting neuropeptides

The R4d ring neurons expressed mRNA for the Dh31 neuropeptide (adjusted p=3.42E-21), while the E-PG neurons expressed mRNA for the Dh31 receptor (adjusted p=0.103). Dh31 has previously been linked to the ellipsoid body and is associated with wake promotion as part of the circadian rhythm (Kunst et al., 2014).

### Electrical isolation of the EB and PB and unique partners within those regions can explain the P-EN2 activity

To probe the source of the differences in observed calcium activity in the P-EN2s between the EB and the PB, we turned to passive cable theory simulations of P-EN2 neurons. We injected current from 3 fictive spikes into Δ7 synapse locations in the PB and observed how potentials propagated along the arbors. Sites within the PB quickly reached equipotential, and voltage losses within the region were minimal. In contrast, sites within the EB experienced only a slight depolarization in response to these current injections, suggesting that the two compartments are electrically isolated (Figure S7O). The P-EN2 activity in the EB may therefore primarily reflect the direct contribution from the P-EGs (Figure S7P) while the activity in the PB would reflect a direct contribution from the Δ7s. The PB P-EN2 activity may only enter the EB when it overcomes some threshold, say when additional current is injected. The P-EN2 activity is tuned to rotational velocities (Green et al., 2017), and thus strong turns may lead to such an additional current source. The super-threshold PB activity would then travel to the EB, leading to a growing shoulder of P-EN2 EB activity at high rotational velocities. Indeed, we see such a shoulder in these conditions (Figure S7Q; see also Figure 3 in Green et al.).

---

**Figure S1 – Related to Figure 1**

A. The offset is the difference between the PVA and the heading. (top) In the dark, error in tracking heading representation activity grows over time and so the offset takes a range of values. (bottom) When a stripe is present, the tracking is accurate over time and so the offset remains relatively fixed.

B. The circular variance for the offset distribution across flies and across conditions.

C. For 4 s sliding windows, the fly’s position (heading) is plotted versus the bump position (PVA). The bump position tracks the stripe position. Data from an example fly is shown here.

D. The slope of the points in the 4 s sliding windows for the fly shown in Figure 1J. The 25^th^ and 75th percentiles are denoted by dotted vertical lines. A slope of 1 is denoted by a solid vertical line.

E. (left) The range between the 25th and 75th percentiles, as taken from the histogram of slopes, for flies in the dark and with a closed-loop stripe present. (right) The median slope in the dark and with a stripe.

**Figure S2 – Related to Figure 2 A.**

A. Comparison between EM skeletons (left) and light level projections (right) for each cell type that was traced. Light level images are maximum intensity projections from confocal imaging stacks. The MultiColor FlpOut (MCFO) technique (Nern et al., 2015) was used to label single neurons. Scale bar is 10 μm.

B. Maximum intensity projections for each genetic line used in this paper. The label in the upper left of each panel is the line name. The label in the upper right refers to the type(s) of experiment that the line was used for.

**Figure S3 – Related to Figure 2**

A. Fraction of identified inputs and outputs for the traced neurons by region. For the Δ7s and ring neurons, where only a few of them were traced in the given regions, relative inputs were extrapolated to what would be expected for a full population. Similarly, as the E-PGs, P-EN1s, P-EGs, and P-EN2s were not traced throughout the input regions for the Δ7s and ring neurons, their total numbers were also extrapolated. Total numbers were estimated by counting cell bodies in light-level confocal stacks of Gal4 lines and are listed next to each type (see Methods for further detail).

**Figure S4 – Related to Figure 3 A.**

A. (top) Mean calcium activity in the ellipsoid body of the E-PGs as recorded with GCaMP6f for flies expressing shi^TS^ in the E-PGs at the permissive temperature when the fly is both in the dark (black) and when a closed-loop stripe is visible (blue). The mean activity during these periods is shown with the horizontal lines. (bottom) The magnitude of fly’s rotational velocity across the trials.

B. As in A, but now at the restrictive temperature, which prevents synaptic vesicle reuptake at E-PG synapses.

C. Cross-correlation coefficient between mean EB E-PG activity and the fly’s rotational velocity across both the light and dark conditions for the permissive (blue) and restrictive (red) temperature trials for the fly shown in A and B.

D. (top) The maximum cross-correlation coefficient (CC) across flies. (bottom) The full width at half maximum (FWHM) for the cross-correlation across flies. Red and blue dots refer to restrictive and permissive trials, respectively.

E. The mean of the overall EB activity in the dark and with a closed-loop (CL) stripe at the permissive temperatures.

F. As above, but at the restrictive temperature.

G. Example changes in ΔF/F around the EB as a fly turns clockwise (blue) and counterclockwise (red) for a variety of stripe positions for four flies.

H. As in Figure 3L, but now with the dots color coded by their PB side and the calcium activity indicator.

I. Calcium activity in the ellipsoid body of the E-PGs as recorded with GCaMP6f for flies expressing shi^TS^ in empty Gal4 at the permissive (top) and restrictive (bottom) temperature. In both conditions, the bump tracks a closed-loop stripe (shown with the blue line).

J. The maximum ΔF/F was found across ROIs for each point in time and the mean of these F values was then calculated over all trials for flies expressing shi^TS^ in empty Gal4 and in the Δ7s. Adjusted p = 0.25, 0.017. 7s

K. As above, but now for the full width at half maximum (FWHM) of the bump of activity. Adjusted p = 0.93, 0.20

L. Fluorescence *in situ* hybridization for GAD+ for the Gall-EB ring neurons measured in (Franconville et al., 2018).

M. The E-PGs and the Δ7s synapse onto the P-EN1s in the protocerebral bridge.

**Figure S5 – Related to Figure 5 A.**

A. Visual inputs were identified through stochastic labeling of broad ring neuron lines (c232 and EB1) with the FLP-out technique. Flies with GCaMP6f in individual neurons were shown an open loop stripe rotating first clockwise and then counterclockwise at 72 °/s to assess visual responses. (top of each panel. The dark period when the stripe disappears behind the fly is indicated with vertical gray bars) Maximum intensity projections of GCaMP6f signals observed in single ring neurons. Multiple different ring neuron morphologies were labeled. Scale bar is 20 μm (bottom of each panel) Visual responses for individual ring neurons. R4d ring neurons consistently showed a strong ipsilateral response.

B. Maximum intensity projection of a population of R4d ring neurons expressing GCaMP7f. The ROIs are denoted by dotted white lines. The central ring and individual glomeruli were analyzed. Scale bar is 20 μm.

C. Mean ring activity for one fly in the ring of the R4d ring neurons, averaged across five trials. The fly is left in the dark for 1 min and then exposed to an open loop stripe rotating first clockwise (blue) and then counterclockwise (magenta) at 72 °/s. The stripe disappears for a 120° arc directly behind the fly, denoted by the gray rectangles. The average activity across the entire dark period is shown with the dotted grey line.

D. The individual clockwise and counterclockwise bouts from C shown on top of one another. The counterclockwise activity is mirrored in time so that that the stripe will be in the same position for the times shown on the x axis.

E. Mean difference in maximum (pink) or minimum intensity (light blue) for the traces shown in D (n = 10).

F. Same as in D, but for the 6 glomeruli.

G. Same as in E, but for the glomeruli (n = 10 flies, ∼6 glomeruli per fly).

H. Calcium activity (green) in the dark for the single neuron. Though this ring neuron is “visual,” it has high baseline activity in the dark. The activity is deconvolved to obtain individual events (orange).

I. Histogram of the inter event intervals (IEI) across trials for an example single ring neuron. The median IEI is noted with the dotted vertical line.

J. Median IEI across all single ring neurons.

**Figure S6 – Related to Figure 5 A.**

A. (top) An example of the E-PG EB activity over time for a control fly. For the first 120 s, the fly is in darkness. For the final 150 s, a closed-loop stripe is shown. The PVA is overlaid in purple. (bottom) The unwrapped PVA, the heading in the dark (magenta), and the stripe position (blue). This example is from the same fly as in Figure 5H, though the fly is now at the permissive temperature.

B. As in Figure S6A, but for the fly shown in Figure 5G, also at the permissive temperature.

**Figure S7 – related to figure 6 A.**

A. Cross-correlation between rotational velocity and the P-EG activity for the fly shown in Figure 6G,H. The fly was subject to 4 different visual conditions: darkness, a closed-loop stripe (CL), a clockwise rotating stripe (CW), and a counterclockwise rotating stripe (CCW).

B. Cross-correlation between rotational velocity and the P-EN2 activity for the fly shown in Figure 6G,H.

C. Maximum correlation coefficient for rotational velocities across flies and across visual conditions. P-EG coefficients are labeled in magenta while P-EN2 coefficients are labeled in green.

D. Maximum correlation coefficient for forward velocities across flies and across visual conditions. P-EG coefficients are labeled in magenta while P-EN2 coefficients are labeled in green.

E. Example calcium activity from a control fly (raised at 21.5° with Gal80ts, UAS-kir transgenes) in the dark (black trace) and with a closed-loop stripe (CL, blue trace).

F. PVA strength vs. rotational velocity in the dark (black) and with a CL stripe (blue) for the fly shown in Figure S7E.

G. Calcium activity from the fly in Figure 6K, now with the CL period shown.

H. As in Figure S7F, now for the fly shown in Figure S7G.

I. The PVA strength vs. rotational velocity in the presence of a closed-loop stripe for control flies (solid) and for flies expressing Kir in the P-EN2s (dotted).

J. The activity in the EB at positions away from the peak activity for the control (dotted) and Kir-expressing P-EN2 (solid) flies.

K. As in Figure S7J, now for the CL trials.

L. Simultaneous calcium activity in the PB for P-EG (magenta, labeled with jRGECO1a) and P-EN2 (green, labeled with GCaMP6f) neurons.

M. The PVA from the right and left PB for the P-EGs (magenta) and P-EN2s (green) vs. the fly’s heading in the dark (black) or the position of a closed-loop stripe (blue).

N. The neuron skeleton for a P-EN2 was imported into NEURON and used in a passive cable model. Three spikes were injected Δ7 at input synapses in the PB and potentials were measured at Δ7 input synapses (in the PB, lower left) and E-PG output synapses (in the EB, lower right). Currents injected into one synapse in the PB rapidly altered the potential at other synapses in the PB. These injected currents had a much smaller impact on the potential in the EB, however, suggesting that the two compartments are electrically isolated. The example traces were generated by activating 30 randomly sampled D7 synapses in a single neuron (PEN2-7R).

O. Mean and standard deviation (shaded) of the maximum potential recorded at each recording synapse in the PB and EB as a function of the number of active input synapses, each of which fires three action potentials. Results were averaged over 10 simulations for each of 8 PEN-2 neurons.

P. The P-EN2s receive input from the P-EGs (blue dotted line) and from the Δ7s (orange dotted line). If these inputs are projected into the EB, the P-EN2 activity will have a hump (black curve). The PB P-EN2 activity may only enter the EB when it overcomes some threshold, say when additional current is injected. P-EN2 activity is tuned to rotational velocities (Green et al., 2017), and thus strong turns may lead to such an additional current source. Suprathreshold PB activity would then travel to the EB, leading to a growing shoulder of P-EN2 EB activity at high rotational velocities.

Q. Measured P-EN2 (green, labeled with GCaMP6f) and E-PG (magenta, labeled with jRGECO1a) activity profiles in the EB. The measured P-EN2 profile is consistent with the modeled profile in P.

**Figure S8: AOTU input to ring neurons is required for flies to maintain arbitrary headings relative to visual cues.**

A. When a closed-loop stripe is presented to the fly (top), the flies tend to keep it in one position across the trial (middle). (bottom) The mean angular position and the mean resultant vector length of the circular distribution is plotted in polar form. The dashed line marks the position where the stripe is directly in front of the fly.

B. Stripe tracking in an Empty Gal4 fly. Two trials are shown, each of which consists of 4 min with a closed-loop stripe. Half-way through the trial, the stripe is moved 120° in open-loop. The periods before and after the stripe moves in open-loop are each shown with different dots, while the two different trials have different shades.

C. The tracking statistics across flies at the permissive (top) and restrictive (bottom) temperature across flies expressing shi^TS^ in empty Gal4 (left), the E-PGs (middle) or the TuBu neurons (right).

D. Stripe tracking behavior for flies expressing shi^TS^ in empty Gal4 or in the Δ7 neurons at the permissive (top) or restrictive (bottom) temperature. The flies are presented with a stripe in closed loop for 2 min.

## References

Amari, S. (1977). Dynamics of pattern formation in lateral-inhibition type neural fields. Biol Cybern 27, 77–87.

Aso, Y., Ray, R.P., Long, X., Bushey, D., Cichewicz, K., Ngo, T.T., Sharp, B., Christoforou, C., Hu, A., Lemire, A.L., et al. (2019). Nitric oxide acts as a cotransmitter in a subset of dopaminergic neurons to diversify memory dynamics. Elife 8.

Ben-Yishai, R., Bar-Or, R.L., and Sompolinsky, H. (1995). Theory of orientation tuning in visual cortex. Proc Natl Acad Sci U S A 92, 3844–3848.

Bender, J.A., Pollack, A.J., and Ritzmann, R.E. (2010). Neural activity in the central complex of the insect brain is linked to locomotor changes. Curr Biol 20, 921–926.

Berens, P. (2009). CircStat: A MATLAB toolbox for circular statistics. J Stat Softw 31, 1–21.

Burak, Y., and Fiete, I.R. (2012). Fundamental limits on persistent activity in networks of noisy neurons. Proc Natl Acad Sci U S A 109, 17645–17650.

Butler, W.N., Smith, K.S., van der Meer, M.A.A., and Taube, J.S. (2017). The Head-Direction Signal Plays a Functional Role as a Neural Compass during Navigation. Curr Biol 27, 1259–1267.

Chaudhuri, R., Gercek, B., Pandey, B., Peyrache, A., and Fiete, I. (2019). The population dynamics of a canonical cognitive circuit. 516021.

Chen, T.W., Wardill, T.J., Sun, Y., Pulver, S.R., Renninger, S.L., Baohan, A., Schreiter, E.R., Kerr, R.A., Orger, M.B., Jayaraman, V., et al. (2013). Ultrasensitive fluorescent proteins for imaging neuronal activity. Nature 499, 295–300.

Clark, B.J., and Taube, J.S. (2012). Vestibular and attractor network basis of the head direction cell signal in subcortical circuits. Front Neural Circuit 6.

Compte, A., Brunel, N., Goldman-Rakic, P.S., and Wang, X.J. (2000). Synaptic mechanisms and network dynamics underlying spatial working memory in a cortical network model. Cereb Cortex 10, 910–923.

Cope, A.J., Sabo, C., Vasilaki, E., Barron, A.B., and Marshall, J.A. (2017). A computational model of the integration of landmarks and motion in the insect central complex. PLoS One 12, e0172325.

Daniels, R.W., Gelfand, M.V., Collins, C.A., and DiAntonio, A. (2008). Visualizing glutamatergic cell bodies and synapses in Drosophila larval and adult CNS. J Comp Neurol 508, 131–152.

Davis, F.P., Nern, A., Picard, S., Reiser, M.B., Rubin, G.M., Eddy, S.R., and Henry, G.L. (2019). A genetic, genomic, and computational resource for exploring neural circuit function. bioRxiv, 385476.

Dobin, A., Davis, C.A., Schlesinger, F., Drenkow, J., Zaleski, C., Jha, S., Batut, P., Chaisson, M., and Gingeras, T.R. (2013). STAR: ultrafast universal RNA-seq aligner. Bioinformatics 29, 15–21.

Donlea, J.M., Pimentel, D., Talbot, C.B., Kempf, A., Omoto, J.J., Hartenstein, V., and Miesenbock, G. (2018). Recurrent Circuitry for Balancing Sleep Need and Sleep. Neuron 97, 378–389 e374.

Dus, M., Ai, M., and Suh, G.S. (2013). Taste-independent nutrient selection is mediated by a brain-specific Na+ /solute co-transporter in Drosophila. Nat Neurosci 16, 526–528.

el Jundi, B., Pfeiffer, K., Heinze, S., and Homberg, U. (2014). Integration of polarization and chromatic cues in the insect sky compass. J Comp Physiol A Neuroethol Sens Neural Behav Physiol 200, 575–589.

el Jundi, B., Warrant, E.J., Byrne, M.J., Khaldy, L., Baird, E., Smolka, J., and Dacke, M. (2015). Neural coding underlying the cue preference for celestial orientation. Proc Natl Acad Sci U S A 112, 11395–11400.

Enell, L., Hamasaka, Y., Kolodziejczyk, A., and Nassel, D.R. (2007). gamma-Aminobutyric acid (GABA) signaling components in Drosophila: immunocytochemical localization of GABA(B) receptors in relation to the GABA(A) receptor subunit RDL and a vesicular GABA transporter. J Comp Neurol 505, 18–31.

Fisher, Y.E., Lu, J., D’Alessandro, I., and Wilson, R.I. (2019). Sensorimotor experience remaps visual input to a heading-direction network. Nature 576, 121–125.

Franconville, R., Beron, C., and Jayaraman, V. (2018). Building a functional connectome of the Drosophila central complex. Elife 7.

Giraldo, Y.M., Leitch, K.J., Ros, I.G., Warren, T.L., Weir, P.T., and Dickinson, M.H. (2018). Sun Navigation Requires Compass Neurons in Drosophila. Curr Biol 28, 2845–2852 e2844.

Golic, K.G., and Lindquist, S. (1989). The FLP recombinase of yeast catalyzes site-specific recombination in the Drosophila genome. Cell 59, 499–509.

Gouwens, N.W., and Wilson, R.I. (2009). Signal propagation in Drosophila central neurons. J Neurosci 29, 6239–6249.

Green, J., Adachi, A., Shah, K.K., Hirokawa, J.D., Magani, P.S., and Maimon, G. (2017). A neural circuit architecture for angular integration in Drosophila. Nature 546, 101–106.

Green, J., Vijayan, V., Mussells Pires, P., Adachi, A., and Maimon, G. (2019). A neural heading estimate is compared with an internal goal to guide oriented navigation. Nat Neurosci 22, 1460–1468.

Guo, P., and Ritzmann, R.E. (2013). Neural activity in the central complex of the cockroach brain is linked to turning behaviors. J Exp Biol 216, 992–1002.

Han, R., Huang, H.-P., Lee, W.-J., Chuang, C.-L., Yen, H.-H., Kao, W.-T., Chang, H.-Y., and Lo, C.-C. (2019). Coordination between stabilizing circuits and updating circuits in spatial orientation working memory. bioRxiv, 819185.

Hanesch, U., Fischbach, K.F., and Heisenberg, M. (1989). Neuronal architecture of the central complex in Drosophila-melanogaster. Cell Tissue Res 257, 343–366.

Hargreaves, E.L., Yoganarasimha, D., and Knierim, J.J. (2007). Cohesiveness of spatial and directional representations recorded from neural ensembles in the anterior thalamus, parasubiculum, medial entorhinal cortex, and hippocampus. Hippocampus 17, 826–841.

Heinze, S., and Homberg, U. (2007). Maplike representation of celestial E-vector orientations in the brain of an insect. Science 315, 995–997.

Heinze, S., and Reppert, S.M. (2011). Sun compass integration of skylight cues in migratory monarch butterflies. Neuron 69, 345–358.

Held, M., Berz, A., Hensgen, R., Muenz, T.S., Scholl, C., Rossler, W., Homberg, U., and Pfeiffer, K. (2016). Microglomerular Synaptic Complexes in the Sky-Compass Network of the Honeybee Connect Parallel Pathways from the Anterior Optic Tubercle to the Central Complex. Front Behav Neurosci 10, 186.

Hines, M.L., Davison, A.P., and Muller, E. (2009). NEURON and Python. Front Neuroinform 3, 1.

Homberg, U., Heinze, S., Pfeiffer, K., Kinoshita, M., and el Jundi, B. (2011). Central neural coding of sky polarization in insects. Philos Trans R Soc Lond B Biol Sci 366, 680–687.

Homberg, U., Hofer, S., Pfeiffer, K., and Gebhardt, S. (2003). Organization and neural connections of the anterior optic tubercle in the brain of the locust, Schistocerca gregaria. Journal of Comparative Neurology 462, 415–430.

Homberg, U., Humberg, T.H., Seyfarth, J., Bode, K., and Perez, M.Q. (2018). GABA immunostaining in the central complex of dicondylian insects. J Comp Neurol 526, 2301–2318.

Homberg, U., and Muller, M. (2016). Ultrastructure of GABA- and Tachykinin-Immunoreactive Neurons in the Lower Division of the Central Body of the Desert Locust. Front Behav Neurosci 10, 230.

Homberg, U., Vitzthum, H., Muller, M., and Binkle, U. (1999). Immunocytochemistry of GABA in the central complex of the locust Schistocerca gregaria: Identification of immunoreactive neurons and colocalization with neuropeptides. Journal of Comparative Neurology 409, 495–507.

Isaacman-Beck, J., Paik, K.C., Wienecke, C.F.R., Yang, H.H., Fisher, Y.E., Wang, I.E., Ishida, I.G., Maimon, G., Wilson, R.I., and Clandinin, T.R. (2019). SPARC: a method to genetically manipulate precise proportions of cells. bioRxiv, 788679.

Itskov, V., Hansel, D., and Tsodyks, M. (2011). Short-Term Facilitation may Stabilize Parametric Working Memory Trace. Front Comput Neurosci 5, 40.

Kahsai, L., Carlsson, M.A., Winther, A.M., and Nassel, D.R. (2012). Distribution of metabotropic receptors of serotonin, dopamine, GABA, glutamate, and short neuropeptide F in the central complex of Drosophila. Neuroscience 208, 11–26.

Kakaria, K.S., and de Bivort, B.L. (2017). Ring Attractor Dynamics Emerge from a Spiking Model of the Entire Protocerebral Bridge. Front Behav Neurosci 11, 8.

Kim, S.S., Hermundstad, A.M., Romani, S., Abbott, L.F., and Jayaraman, V. (2019). Generation of stable heading representations in diverse visual scenes. Nature 576, 126–131.

Kim, S.S., Rouault, H., Druckmann, S., and Jayaraman, V. (2017). Ring attractor dynamics in the Drosophila central brain. Science 356, 849–853.

Kitamoto, T. (2002). Conditional disruption of synaptic transmission induces male-male courtship behavior in Drosophila. Proc Natl Acad Sci U S A 99, 13232–13237.

Knierim, J.J., Kudrimoti, H.S., and McNaughton, B.L. (1998). Interactions between idiothetic cues and external landmarks in the control of place cells and head direction cells. J Neurophysiol 80, 425–446.

Knierim, J.J., and Zhang, K.C. (2012). Attractor dynamics of spatially correlated neural activity in the limbic system. Annual Review of Neuroscience, Vol 35 35, 267–285.

Krause, T., Spindler, L., Poeck, B., and Strauss, R. (2019). Drosophila Acquires a Long-Lasting Body-Size Memory from Visual Feedback. Curr Biol 29, 1833–1841 e1833.

Kunst, M., Hughes, M.E., Raccuglia, D., Felix, M., Li, M., Barnett, G., Duah, J., and Nitabach, M.N. (2014). Calcitonin gene-related peptide neurons mediate sleep-specific circadian output in Drosophila. Curr Biol 24, 2652–2664.

Kuntz, S., Poeck, B., and Strauss, R. (2017). Visual Working Memory Requires Permissive and Instructive NO/cGMP Signaling at Presynapses in the Drosophila Central Brain. Curr Biol 27, 613–623.

Lee, D., Huang, T.H., De La Cruz, A., Callejas, A., and Lois, C. (2017). Methods to investigate the structure and connectivity of the nervous system. Fly (Austin) 11, 224–238.

Lee, D., and O’Dowd, D.K. (1999). Fast excitatory synaptic transmission mediated by nicotinic acetylcholine receptors in Drosophila neurons. J Neurosci 19, 5311–5321.

Li, B., and Dewey, C.N. (2011). RSEM: accurate transcript quantification from RNA-Seq data with or without a reference genome. BMC Bioinformatics 12, 323.

Liang, X., Ho, M.C.W., Zhang, Y., Li, Y., Wu, M.N., Holy, T.E., and Taghert, P.H. (2019). Morning and Evening Circadian Pacemakers Independently Drive Premotor Centers via a Specific Dopamine Relay. Neuron 102, 843–857 e844.

Lin, C.Y., Chuang, C.C., Hua, T.E., Chen, C.C., Dickson, B.J., Greenspan, R.J., and Chiang, A.S. (2013). A comprehensive wiring diagram of the protocerebral bridge for visual information processing in the Drosophila brain. Cell reports 3, 1739–1753.

Liu, G., Seiler, H., Wen, A., Zars, T., Ito, K., Wolf, R., Heisenberg, M., and Liu, L. (2006). Distinct memory traces for two visual features in the Drosophila brain. Nature 439, 551–556.

Liu, S., Liu, Q., Tabuchi, M., and Wu, M.N. (2016). Sleep Drive Is Encoded by Neural Plastic Changes in a Dedicated Circuit. Cell 165, 1347–1360.

Liu, W.W., and Wilson, R.I. (2013). Glutamate is an inhibitory neurotransmitter in the Drosophila olfactory system. Proc Natl Acad Sci U S A 110, 10294–10299.

Long, X., Colonell, J., Wong, A.M., Singer, R.H., and Lionnet, T. (2017). Quantitative mRNA imaging throughout the entire Drosophila brain. Nat Methods 14, 703–706.

Martin, M. (2011). Cutadapt removes adapter sequences from high-throughput sequencing reads. 2011 17, 3.

Martin-Pena, A., Acebes, A., Rodriguez, J.R., Chevalier, V., Casas-Tinto, S., Triphan, T., Strauss, R., and Ferrus, A. (2014). Cell types and coincident synapses in the ellipsoid body of Drosophila. European Journal of Neuroscience 39, 1586–1601.

Meissner, G.W., Nern, A., Singer, R.H., Wong, A.M., Malkesman, O., and Long, X. (2019). Mapping Neurotransmitter Identity in the Whole-Mount Drosophila Brain Using Multiplex High-Throughput Fluorescence in Situ Hybridization. Genetics 211, 473–482.

Muir, G.M., Brown, J.E., Carey, J.P., Hirvonen, T.P., Della Santina, C.C., Minor, L.B., and Taube, J.S. (2009). Disruption of the Head Direction Cell Signal after Occlusion of the Semicircular Canals in the Freely Moving Chinchilla. Journal of Neuroscience 29, 14521–14533.

Nern, A., Pfeiffer, B.D., and Rubin, G.M. (2015). Optimized tools for multicolor stochastic labeling reveal diverse stereotyped cell arrangements in the fly visual system. Proc Natl Acad Sci U S A 112, E2967–2976.

Neuser, K., Triphan, T., Mronz, M., Poeck, B., and Strauss, R. (2008). Analysis of a spatial orientation memory in Drosophila. Nature 453, 1244–1247.

Ocko, S.A., Hardcastle, K., Giocomo, L.M., and Ganguli, S. (2018). Emergent elasticity in the neural code for space. Proceedings of the National Academy of Sciences.

Ofstad, T.A., Zuker, C.S., and Reiser, M.B. (2011). Visual place learning in Drosophila melanogaster. Nature 474, 204–207.

Omoto, J.J., Keles, M.F., Nguyen, B.M., Bolanos, C., Lovick, J.K., Frye, M.A., and Hartenstein, V. (2017). Visual Input to the Drosophila Central Complex by Developmentally and Functionally Distinct Neuronal Populations. Curr Biol 27, 1098–1110.

Omoto, J.J., Nguyen, B.M., Kandimalla, P., Lovick, J.K., Donlea, J.M., and Hartenstein, V. (2018). Neuronal Constituents and Putative Interactions Within the Drosophila Ellipsoid Body Neuropil. Front Neural Circuits 12, 103.

Page, H.J.I., and Jeffery, K.J. (2018). Landmark-Based Updating of the Head Direction System by Retrosplenial Cortex: A Computational Model. Front Cell Neurosci 12, 191.

Park, J.Y., Dus, M., Kim, S., Abu, F., Kanai, M.I., Rudy, B., and Suh, G.S.B. (2016). Drosophila SLC5A11 Mediates Hunger by Regulating K(+) Channel Activity. Curr Biol 26, 1965–1974.

Peyrache, A., Lacroix, M.M., Petersen, P.C., and Buzsaki, G. (2015). Internally organized mechanisms of the head direction sense. Nat Neurosci.

Phillips-Portillo, J. (2012). The central complex of the flesh fly, Neobellieria bullata: recordings and morphologies of protocerebral inputs and small-field neurons. J Comp Neurol 520, 3088–3104.

Renart, A., Song, P., and Wang, X.J. (2003). Robust spatial working memory through homeostatic synaptic scaling in heterogeneous cortical networks. Neuron 38, 473–485.

Schlegel, P., Texada, M.J., Miroschnikow, A., Schoofs, A., Huckesfeld, S., Peters, M., Schneider-Mizell, C.M., Lacin, H., Li, F., Fetter, R.D., et al. (2016). Synaptic transmission parallels neuromodulation in a central food-intake circuit. Elife 5.

Seeholzer, A., Deger, M., and Gerstner, W. (2019). Stability of working memory in continuous attractor networks under the control of short-term plasticity. PLoS computational biology 15, e1006928.

Seelig, J.D., Chiappe, M.E., Lott, G.K., Dutta, A., Osborne, J.E., Reiser, M.B., and Jayaraman, V. (2010). Two-photon calcium imaging from head-fixed Drosophila during optomotor walking behavior. Nat Methods 7, 535–540.

Seelig, J.D., and Jayaraman, V. (2013). Feature detection and orientation tuning in the Drosophila central complex. Nature 503, 262–266.

Seelig, J.D., and Jayaraman, V. (2015). Neural dynamics for landmark orientation and angular path integration. Nature 521, 186–191.

Shiozaki, H.M., and Kazama, H. (2017). Parallel encoding of recent visual experience and self-motion during navigation in Drosophila. Nat Neurosci 20, 1395–1403.

Skaggs, W.E., Knierim, J.J., Kudrimoti, H.S., and McNaughton, B.L. (1995). A model of the neural basis of the rat’s sense of direction. Adv Neural Inf Process Syst 7, 173–180.

Soumillon, M., Cacchiarelli, D., Semrau, S., van Oudenaarden, A., and Mikkelsen, T.S. (2014). Characterization of directed differentiation by high-throughput single-cell RNA-Seq. bioRxiv, 003236.

Stone, T., Webb, B., Adden, A., Weddig, N.B., Honkanen, A., Templin, R., Wcislo, W., Scimeca, L., Warrant, E., and Heinze, S. (2017). An Anatomically Constrained Model for Path Integration in the Bee Brain. Curr Biol.

Su, T.S., Lee, W.J., Huang, Y.C., Wang, C.T., and Lo, C.C. (2017). Coupled symmetric and asymmetric circuits underlying spatial orientation in fruit flies. Nat Commun 8, 139.

Sun, Y., Nern, A., Franconville, R., Dana, H., Schreiter, E.R., Looger, L.L., Svoboda, K., Kim, D.S., Hermundstad, A.M., and Jayaraman, V. (2017). Neural signatures of dynamic stimulus selection in Drosophila. Nat Neurosci 20, 1104–1113.

Talay, M., Richman, E.B., Snell, N.J., Hartmann, G.G., Fisher, J.D., Sorkac, A., Santoyo, J.F., Chou-Freed, C., Nair, N., Johnson, M., et al. (2017). Transsynaptic Mapping of Second-Order Taste Neurons in Flies by trans-Tango. Neuron 96, 783–795 e784.

Taube, J.S. (2007). The head direction signal: Origins and sensory-motor integration. Annual Review of Neuroscience 30, 181–207.

Taube, J.S., Muller, R.U., and Ranck, J.B. (1990a). Head-direction cells recorded from the postsubiculum in freely moving rats. 1. Description and quantitative analysis. Journal of Neuroscience 10, 420–435.

Taube, J.S., Muller, R.U., and Ranck, J.B. (1990b). Head-direction cells recorded from the postsubiculum in freely moving rats. 2. Effects of environmental manipulations. Journal of Neuroscience 10, 436–447.

Triphan, T., Poeck, B., Neuser, K., and Strauss, R. (2010). Visual targeting of motor actions in climbing Drosophila. Curr Biol 20, 663–668.

Turner-Evans, D., Wegener, S., Rouault, H., Franconville, R., Wolff, T., Seelig, J.D., Druckmann, S., and Jayaraman, V. (2017). Angular velocity integration in a fly heading circuit. Elife 6.

Valerio, S., and Taube, J.S. (2012). Path integration: how the head direction signal maintains and corrects spatial orientation. Nature Neuroscience 15, 1445–1453.

Varga, A.G., and Ritzmann, R.E. (2016). Cellular Basis of Head Direction and Contextual Cues in the Insect Brain. Curr Biol 26, 1816–1828.

Vitzthum, H., Muller, M., and Homberg, U. (2002). Neurons of the Central Complex of the Locust Schistocerca gregaria are Sensitive to Polarized Light. J Neurosci 22, 1114–1125.

Vogelstein, J.T., Packer, A.M., Machado, T.A., Sippy, T., Babadi, B., Yuste, R., and Paninski, L. (2010). Fast nonnegative deconvolution for spike train inference from population calcium imaging. J Neurophysiol 104, 3691–3704.

Wolff, T., Iyer, N.A., and Rubin, G.M. (2015). Neuroarchitecture and neuroanatomy of the Drosophila central complex: A GAL4-based dissection of protocerebral bridge neurons and circuits. J Comp Neurol 523, 997–1037.

Wolff, T., and Rubin, G.M. (2018). Neuroarchitecture of the Drosophila central complex: A catalog of nodulus and asymmetrical body neurons and a revision of the protocerebral bridge catalog. J Comp Neurol 526, 2585–2611.

Wu, S., and Amari, S. (2005). Computing with continuous attractors: stability and online aspects. Neural Comput 17, 2215–2239.

Wu, S., Hamaguchi, K., and Amari, S. (2008). Dynamics and computation of continuous attractors. Neural Comput 20, 994–1025.

Xia, S., Miyashita, T., Fu, T.F., Lin, W.Y., Wu, C.L., Pyzocha, L., Lin, I.R., Saitoe, M., Tully, T., and Chiang, A.S. (2005). NMDA receptors mediate olfactory learning and memory in Drosophila. Curr Biol 15, 603–615.

Xie, X., Hahnloser, R.H., and Seung, H.S. (2002). Double-ring network model of the head-direction system. Phys Rev E Stat Nonlin Soft Matter Phys 66, 041902.

Xie, X., Tabuchi, M., Brown, M.P., Mitchell, S.P., Wu, M.N., and Kolodkin, A.L. (2017). The laminar organization of the Drosophila ellipsoid body is semaphorin-dependent and prevents the formation of ectopic synaptic connections. Elife 6.

Yoganarasimha, D., Yu, X.T., and Knierim, J.J. (2006). Head direction cell representations maintain internal coherence during conflicting proximal and distal cue rotations: Comparison with hippocampal place cells. Journal of Neuroscience 26, 622–631.

Young, J.M., and Armstrong, J.D. (2010). Structure of the adult central complex in Drosophila: Organization of distinct neuronal subsets. Journal of Comparative Neurology 518, 1500–1524.

Zhang, K. (1996). Representation of spatial orientation by the intrinsic dynamics of the head-direction cell ensemble: A theory. Journal of Neuroscience 16, 2112–2126.

Zhang, W., Basile, A.S., Gomeza, J., Volpicelli, L.A., Levey, A.I., and Wess, J. (2002). Characterization of central inhibitory muscarinic autoreceptors by the use of muscarinic acetylcholine receptor knock-out mice. J Neurosci 22, 1709–1717.

Zhang, Z., Li, X., Guo, J., Li, Y., and Guo, A. (2013). Two clusters of GABAergic ellipsoid body neurons modulate olfactory labile memory in Drosophila. J Neurosci 33, 5175–5181.

Zheng, Z., Lauritzen, J.S., Perlman, E., Robinson, C.G., Nichols, M., Milkie, D., Torrens, O., Price, J., Fisher, C.B., Sharifi, N., et al. (2018). A Complete Electron Microscopy Volume of the Brain of Adult Drosophila melanogaster. Cell 174, 730–743 e722.

